# A CRISPRi/a platform in iPSC-derived microglia uncovers regulators of disease states

**DOI:** 10.1101/2021.06.16.448639

**Authors:** Nina M. Dräger, Sydney M. Sattler, Cindy Tzu-Ling Huang, Olivia M. Teter, Kun Leng, Sayed Hadi Hashemi, Jason Hong, Giovanni Aviles, Claire D. Clelland, Lihong Zhan, Joe C. Udeochu, Lay Kodama, Andrew B. Singleton, Mike A. Nalls, Justin Ichida, Michael E. Ward, Faraz Faghri, Li Gan, Martin Kampmann

**Affiliations:** Institute for Neurodegenerative Diseases, University of California, San Francisco, San Francisco, CA, USA; Gladstone Institute of Neurological Disease, San Francisco, CA, USA; UC Berkeley-UCSF Graduate Program in Bioengineering, University of California San Francisco, San Francisco, CA, USA; Biomedical Sciences Graduate Program, University of California, San Francisco, San Francisco, CA, USA; Medical Scientist Training Program, University of California, San Francisco, San Francisco, CA, USA; Department of Computer Science, University of Illinois at Urbana-Champaign, Urbana, IL, USA; Department of Neurology, University of California, San Francisco, CA, 94158, USA.; Neuroscience Graduate Program, University of California, San Francisco, San Francisco, CA, USA; Center for Alzheimer’s and Related Dementias, National Institutes of Health, Bethesda, MD, USA; Laboratory of Neurogenetics, National Institute on Aging, National Institutes of Health, Bethesda, MD, USA; Data Tecnica International, LLC, Glen Echo, MD, USA; Department of Stem Cell Biology and Regenerative Medicine, Keck School of Medicine, University of Southern California, Los Angeles, CA, USA; Eli and Edythe Broad CIRM Center for Regenerative Medicine and Stem Cell Research at USC, Los Angeles, CA, USA; Zilkha Neurogenetic Institute, Keck School of Medicine, University of Southern California, Los Angeles, CA, USA; National Institute of Neurological Disorders and Stroke, National Institutes of Health, Bethesda, MD, USA; Helen and Robert Appel Alzheimer’s Disease Research Institute, Brain and Mind Research Institute, Weill Cornell Medicine, New York, NY, USA; Department of Biochemistry and Biophysics, University of California, San Francisco, San Francisco, CA, USA; Chan Zuckerberg Biohub, San Francisco, CA, USA

## Abstract

Microglia are emerging as key drivers of neurological diseases. However, we lack a systematic understanding of the underlying mechanisms. Here, we present a screening platform to systematically elucidate functional consequences of genetic perturbations in human iPSC-derived microglia. We developed an efficient eight-day protocol for the generation of microglia-like cells based on the inducible expression of six transcription factors. We established inducible CRISPR interference and activation in this system and conducted three screens targeting the “druggable genome”. These screens uncovered genes controlling microglia survival, activation and phagocytosis, including neurodegeneration-associated genes. A screen with single-cell RNA sequencing as the readout revealed that these microglia adopt a spectrum of states mirroring those observed in human brains and identified regulators of these states. A disease-associated state characterized by SPP1 expression was selectively depleted by CSF1R inhibition. Thus, our platform can systematically uncover regulators of microglia states, enabling their functional characterization and therapeutic targeting.

## INTRODUCTION

Historically, neuroscience has investigated brain function and disease through a neuron-centric lens, relegating glia to the sidelines. Neuroinflammation has typically been viewed as a secondary, reactive aspect of disease. More recently, however, key roles have emerged for glial cell types, including microglia, the innate immune cells of the brain. It is now widely accepted that microglia have a central role in brain development and homeostasis as well as in the pathogenesis of many brain disorders^1^. Over the last decade, human genetics have pointed to a central role for microglia in brain diseases such as Alzheimer’s Disease (AD)^2^, where specific disease-associated genetic variants likely act in microglia, redefining them as potential drivers of AD. To understand the molecular mechanisms underlying the role of microglia in disease and target them therapeutically, it is necessary to bridge the gap between disease-associated genetic variants and changes in microglial function.

A major challenge is that microglia adopt a large number of distinct functional states in health and disease. In homeostatic states, microglia survey their local environment, phagocytose myelin and cell debris, and monitor neuronal activity^3^. In disease states, microglia can play beneficial roles, but they are also responsible for an increased production of proinflammatory cytokines, an exacerbated inflammatory response and secrete toxic factors to directly or indirectly damage neurons^4, 5^. Eventually, microglia exhibit pathological features, such as mitochondrial and endolysosomal dysfunction, impaired phagocytosis, and increased production of reactive oxygen species (ROS)^6^.

Microglial states in health and disease are actively being mapped on the molecular level in mice and humans^7–13^. However, we do not systematically understand how these distinct microglial states contribute to brain function and disease, or the molecular mechanisms regulating these states.

A promising approach to tackle these questions is enabled by CRISPR-based functional genomics in differentiated human cell types^14^. Pooled CRISPR interference (CRISPRi) and CRISPR activation (CRISPRa) screens enable scalable modeling of changes in gene expression and genetic screens to uncover regulatory mechanisms. When combined with induced pluripotent stem cell (iPSC) technology, they enable the investigation of cell-type specific biology in human cells, including those derived from patients^14^. We recently provided a proof of principle for this strategy by establishing CRISPRi and CRISPRa platforms for genetic screens in iPSC-derived neurons^15, 16^. However, such screens have not previously been implemented in iPSC-derived microglia due to challenges inherent in available differentiation protocols. Pooled CRISPR screens rely on lentiviral transduction to introduce libraries of single guide RNAs (sgRNAs), but mature microglia are difficult to transduce with lentivirus. This problem could be overcome by introducing sgRNAs at the iPSC stage. However, most existing protocols are lengthy and aim to recapitulate human microglia ontogeny^17–23^, resulting in population bottlenecks during differentiation, which can skew the representation of the sgRNA library.

To overcome these challenges, we developed a different approach for the generation of iPSC-derived microglia by generating a human iPSC line inducibly expressing six transcription factors that enable the generation of microglia-like cells in a rapid and efficient eight-day protocol. These induced-transcription factor microglia-like cells (iTF-Microglia) resemble other iPSC-derived microglia in their expression profiles, response to inflammatory stimuli, phagocytic capabilities, and capacity to be co-cultured with iPSC-derived neurons^17–23^. By integrating inducible CRISPRi/a machinery into this cell line, we developed a genetic screening system that enables robust knockdown and overexpression of endogenous genes in human microglia. Using this platform, we conducted pooled CRISPRi and CRISPRa screens for modifiers of survival, phagocytosis and inflammatory activation, which uncovered microglia-specific genes controlling these phenotypes. A screen with single-cell RNA sequencing as the readout revealed that these microglia adopt a spectrum of states mirroring those observed in human brains, and pinpointed regulators of specific states, which can enable the functional characterization and therapeutic targeting of these states.

## RESULTS

### Inducible transcription factors enable rapid and scalable production of microglia-like cells (iTF-Microglia)

High-throughput genetic CRISPRi/a screens are a powerful discovery tool in human iPSC-derived neurons^15, 16^. However, such screens have not yet been conducted in iPSC-derived microglia. One obstacle has been the fact that until very recently^24^, existing differentiation protocols involved a long, multi-step procedure, which recapitulates the human microglia ontogeny. We set out to create a fast, robust and scalable differentiation protocol to differentiate iPSCs to microglia-like cells for use in CRISPR screens. To this end, we developed a strategy based on direct cell fate conversion by overexpression of microglia fate-determining transcription factors.

Based on transcriptomic and developmental data^25–27^, we selected six transcription factors highly expressed in human microglia. PU.1 and interferon regulatory factor-8 (IRF-8) are known to be crucial for microgliogenesis^28^. MAFB increases during microglia development and promotes an anti-inflammatory phenotype^29^, while CEBP*α* and CEBP*β*^30^ and IRF-5^31^ regulate a wide range of inflammatory mediators. We engineered an iPSC line with two integrated cassettes of three transcription factors each. Cassettes for the doxycycline-inducible expression of transgenic PU.1/CEBP*β*/IRF5 and MAFB/CEBP*α*/IRF8 were integrated into the CLYBL and AAVS1 safe harbor loci, respectively, in the WTC11 iPSC line (Fig. 1a).

**Figure 1:**
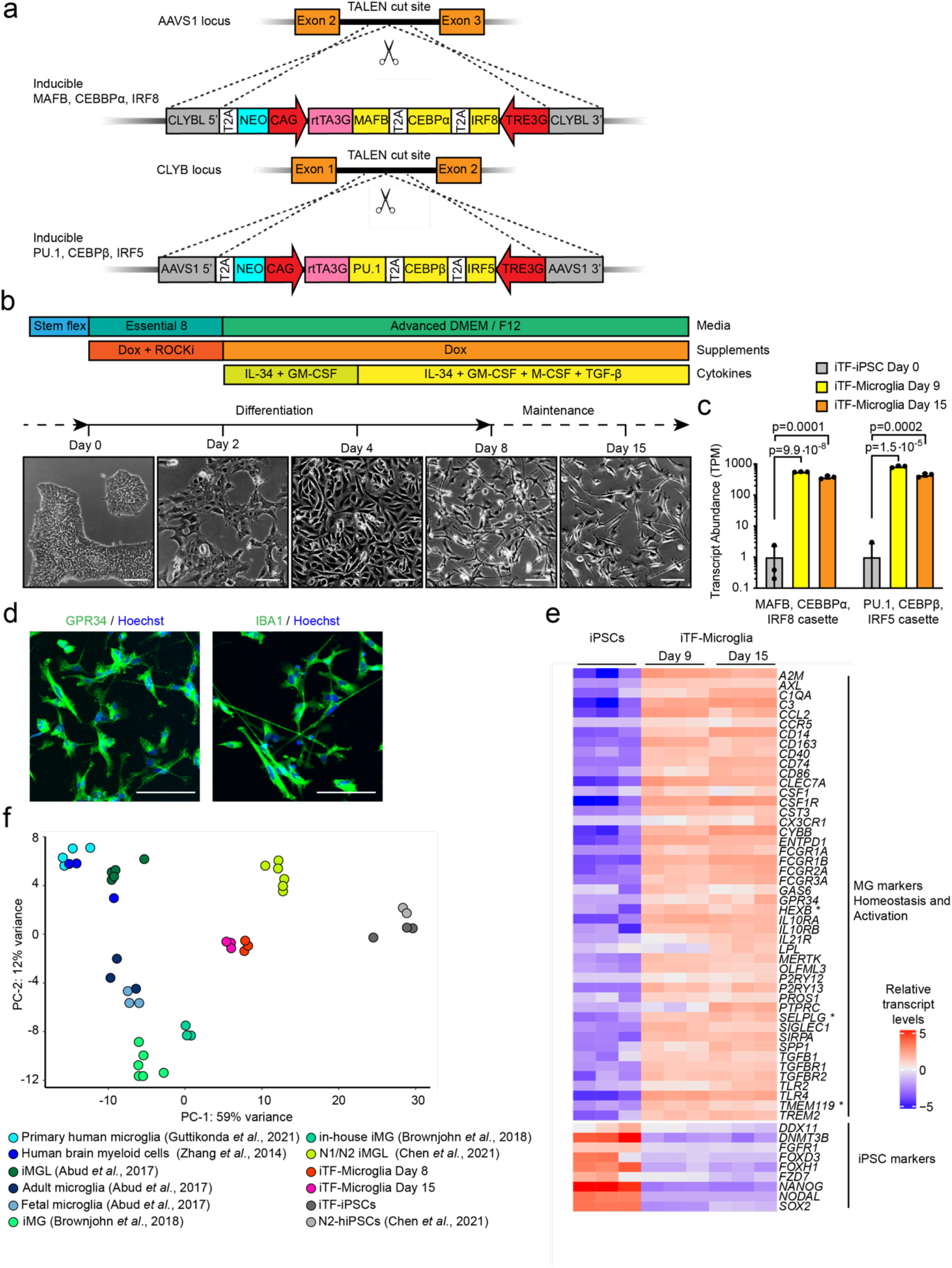
Rapid differentiation of iPSCs into microglia-like cells (iTF-Microglia) by transcription factor induction. **a,** Strategy for stable integration of six transcription factors integrated in AAVS1 and CLYBL loci by TALEN-mediated integration: The doxycycline-inducible reverse transcriptional activator (rtTA3G) is driven by the constitutive CAG promoter. Human MAFB, CEBPα and IRF8 are driven by the tet response element (TRE3G) in the AAVS1 locus. Human PU.1, CEBP*β* and IRF5 are driven by TRE3G in the CLYBL locus. All transcription factors are separated from each other via T2A ribosome skipping sequences. **b,** Overview of the differentiation process for generating iTF-Microglia. *Top*: timeline with media and cytokines, *bottom*: representative phase-contrast images of cells on the indicated days. Scale bar: 100 μm. **c,** Expression of six inducible transcription factors during iTF-Microglia differentiation. Transcript abundance (TPM) of MAFB, CEBPα, IRF8 cassette and the PU.1, CEBPβ, IRF5 cassette at Day 0, Day 9 and Day 15 of differentiation. n = 3 biological replicates, p values from two-tailed Student’s t-test. **d,** Representative immunofluorescence micrographs of iTF-Microglia on Day 8 of differentiation stained for microglia markers GPR34 and IBA1. Nuclei were labeled by Hoechst 33342. Scale bar: 100 μm. **e,** Expression of iPSC and microglia marker genes in iPSCs and derived iTF-Microglia on Day 9 and Day 15 of differentiation. The heatmap displays normalized and gene-centered transcripts per million (TPM) counts for selected genes (rows) for 3 biological replicates of timepoints (columns).iTF-Microglia express microglia homeostatic markers and activation markers, while losing their expression of iPSC markers. Asterisks highlight microglia-selective markers. **f,** Principal component analysis (PCA) on the expression of microglia marker genes of iTF-Microglia, human adult ex-vivo microglia^93^, fetal and adult microglia^17^, human myeloid cells^25^, other iPSC-microglia^17, 22, 24^ and iPSCs (this study and ref. ^24^). Each dot reflects an independent biological sample. Colors represent the different cell types.

We established a simple three-step protocol to differentiate these iPSCs into microglia-like cells, which we will refer to as iTF-Microglia, in only 8 days (Fig. 1b). After doxycycline induction of transcription factor expression on Day 0, media was supplemented with cytokines GM-CSF and IL-34 on Day 2 to promote differentiation and survival. On Day 4, the media was additionally supplemented with the cytokines M-CSF and TGF-*β*. iTF-Microglia reached a fully ramified morphology on Day 8 and maintained excellent viability for at least another 8 days (Fig. 1b). We generally have continued doxycycline supplementation beyond Day 8; however, this is not necessary for survival (Extended Data Fig. 1a,b). We confirmed robust inducible expression of the transgenic transcription factors (Fig. 1c).

The canonical microglia markers GPR34 and IBA1 were expressed in the iTF-Microglia at Day 8 of differentiation (Fig. 1d). To confirm the cell type identity of iTF-Microglia, we conducted RNA-Seq and compared transcript levels of iPSC markers and microglia markers in Day-9 and Day-15 iTF-iMicroglia to the parental iPSCs (Fig. 1e, Supplementary Table 1). As expected, the expression of iPSC markers was drastically reduced in iTF-Microglia at Day 9 and Day 15, whereas microglia markers were induced. Some markers, such as *P2RY12*, *CSF1R*, *CYBB* and *CD14* slightly increased their expression from Day 9 to Day 15, indicating further incremental maturation from Day 9 to Day 15. While the transcriptomic signature of our microglia was distinct from primary human microglia (Fig. 1f, Extended Data Fig. 1c), it was comparable to that of several other iPSC-derived microglia protocols.

In conclusion, our transcriptomic and immunofluorescence results indicate robust expression of microglia markers in iTF-Microglia. Importantly, our novel differentiation strategy is compatible with large-scale pooled sgRNA screens, whereas classical protocols create population bottlenecks (Extended Data Fig. 1d).

### Functional characterization of iTF-Microglia

Next, we asked whether iTF-Microglia recapitulated cellular functions of human microglia. Microglia are the professional phagocytes in the brain, enabling them to clear neuronal debris, prune synapses and engulf pathogens. Using live-cell imaging and flow cytometry, we found that iTF-Microglia robustly phagocytose fluorescent beads (Extended Data Fig. 2a) and rat synaptosomes labeled with the pH-sensitive fluorescent dye pHrodo (Fig. 2a, b, Extended Data Fig. 2b). As expected, phagocytosis could be attenuated by the actin polymerization inhibitor Cytochalasin D, since phagocytosis depends on actin dynamics (Fig. 2a, b, Extended Data Fig. 2c).

**Figure 2:**
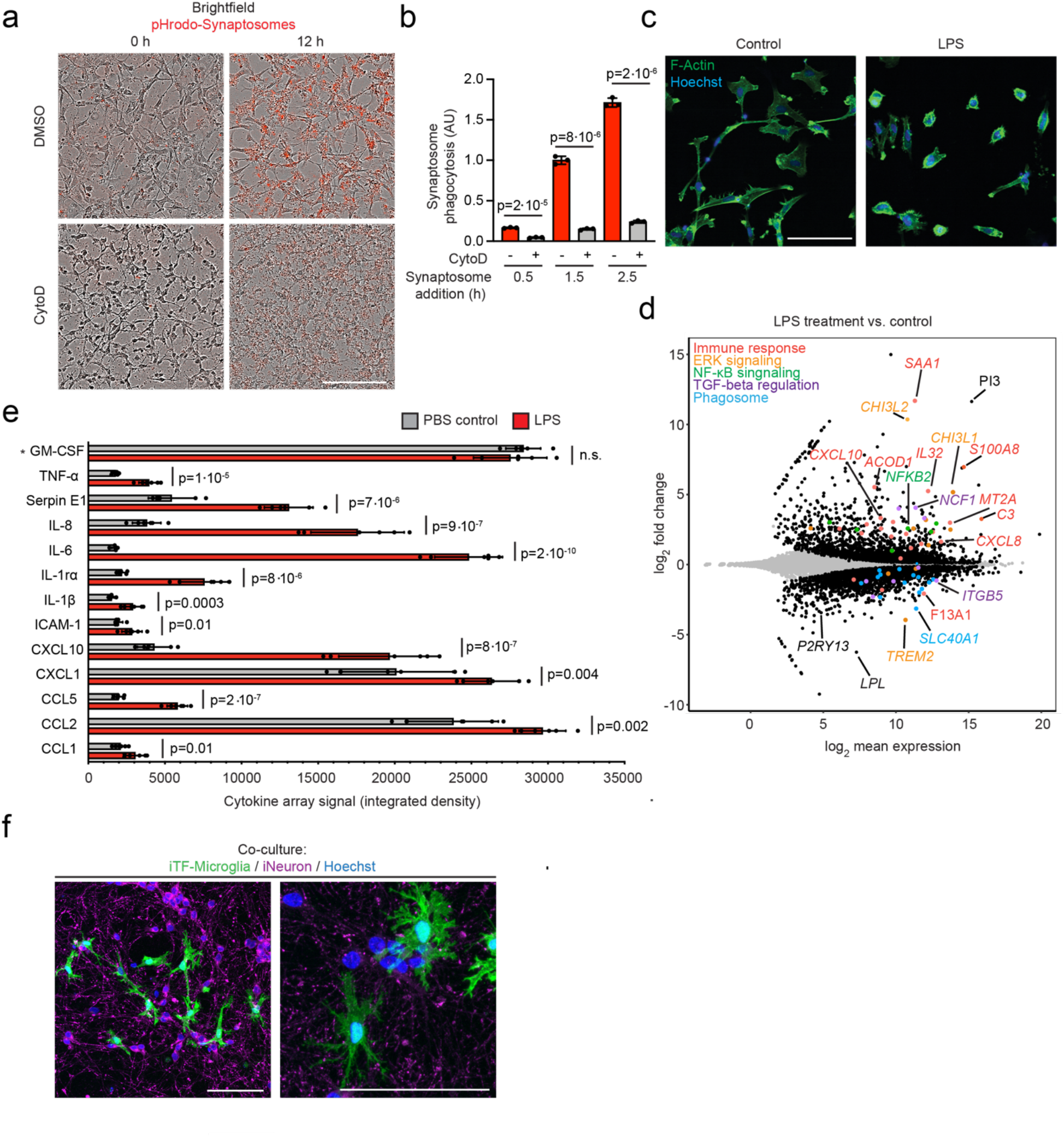
Functional characterization of iTF-Microglia. **a,** Phagocytosis of pHrodo-red-labeled rat brain-derived synaptosomes by iTF-Microglia. Representative images at 0 h and 12 h after synaptosome addition are shown. Treatment with 5 μM actin polymerization inhibitor Cytochalasin D decreases phagocytosis. Scale bar: 100 μm. **b,** Phagocytosis of pHrodo-labeled rat brain-derived synaptosomes with or without Cytochalasin D treatment was quantified by flow cytometry at 0.5 h, 1.5 h and 2.5 h after synaptosome addition (mean +/- sd, n = three biological replicates; p values from two-tailed Student’s t-test). **c,** Morphological changes of iTF-Microglia after LPS treatment are visualized by fluorescence microscopy. Samples were treated for 24 h with 100 ng/ml LPS or buffer control and fixed samples were stained with AlexaFluor 488-phalloidin for F-actin (green) and with Hoechst 33342 for nuclei (blue). Scale bar: 100 μm. **d,** Transcriptomic changes caused by 50 ng/ml lipo-polysaccharide (LPS) treatment in Day 15 iTF-Microglia (n = three biological replicates). Differentially expressed genes (padj < 0.05) are labeled in black (increase). Other colors label genes associated with specific pathways that are discussed in the main text. **e,** Cytokines secreted by iTF-Microglia. Analysis of cytokine array signal (integrated density of dot blots) from supernatants of cultures treated with LPS or buffer control (mean +/- sd, n = 6 biological replicates; p values from two-tailed Student’s t-test). Asterisk: GM-CSF is a component of the culture media. **f,** Co-culture with iPSC-derived excitatory neurons promotes ramified morphology of iTF-Microglia. Representative fluorescence micrographs at low and high magnification of Day 9 iTF-Microglia after 24 hours in co-culture. iTF-Microglia express membrane-localized Lck-mNeonGreen (green). Neurons are stained for the pre-synaptic marker synaptophysin (magenta). Nuclei are stained with Hoechst 33342 (blue). Scale bars = 100 µm.

Microglia express pattern recognition receptors such as Toll-like receptors that mediate the inflammatory response to pathogen-associated patterns including bacterial-derived lipopolysaccharide (LPS). To test the inflammatory response of iTF-Microglia, we stimulated them with LPS for 24 h and evaluated morphological changes after staining for F-actin. LPS-stimulated iTF-Microglia were less ramified, and instead displayed the ameboid morphology characteristic of activated microglia (Fig. 2c, Extended Data Fig. 2d). In addition to the observed morphological changes, we examined transcriptomic alterations after LPS challenge by RNA-Seq (Fig. 2d, Supplementary Table 2). As anticipated, many of the highly upregulated genes were immune response genes such as *C3*, *CXCL10*, *IL32* and *SAA1*. Moreover, several upregulated genes were members of the NF-*κ*B pathway. Downregulated genes included *TREM2*, markers of homeostatic microglia, such as *P2RY13*, and members of the TGF-*β* signaling pathway, such as *SLC40A1*. Transcriptomic changes in response to LPS were substantially overlapping with those observed in iPSC-derived microglia we generated following an alternative, previously published^22^ protocol (Extended Data Fig.2e, Supplementary Table 2).

To examine cytokine secretion of iTF-Microglia, we measured the abundance of 36 cytokines secreted in standard culture conditions or following LPS stimulation. Control buffer-treated iTF-Microglia secreted most cytokines at low levels, but higher levels of CCL2 and CXCL1, suggesting the presence of activated cells under control conditions (Fig. 2e), consistent with previous reports suggesting that even primary microglia become partially activated when cultured^32^. When stimulated with LPS, levels of most secreted cytokines increased; the most increased cytokine levels where IL-6 with a 14-fold increase and IL-8 and CXCL10, both increased over 4-fold (Fig. 2e).

During human development, microglia precursors enter the developing brain and mature together with neurons into fully functional microglia. To test if neurons can promote iTF-Microglia maturation, we co-cultured iTF-Microglia with iPSC-derived glutamatergic neurons (iNeurons) in medium optimized for survival and functionality of both cell types (see Methods for details). Day 8 iTF-Microglia expressing GFP differentiated in mono-culture were co-cultured with iNeurons for one week. Remarkably, the co-cultured iTF-Microglia displayed a pronounced ramified morphology (Fig. 2f). In conclusion, we show that iTF-Microglia effectively phagocytose synaptosomes, respond to LPS and can be co-cultured with iPSC-derived neurons.

### Durable gene knockdown and overexpression by CRISPRi and CRISPRa in iTF-Microglia

Next, we established CRISPRi and CRISPRa in iTF-Microglia to enable robust knockdown and overexpression of endogenous genes, as well as large-scale loss- and gain-of-function genetic screens. Following the strategy, we previously established in human iPSC-derived neurons^15, 16^, we stably integrated constitutive CRISPRi machinery, inducible CRISPRi machinery, or inducible CRISPRa machinery into safe-harbor loci of iPSCs also engineered with the inducible microglial transcription factors (Fig. 3a). We confirmed a normal karyotype for the resulting monoclonal cell lines (Extended Data Fig. 3). Inducible CRISPRi/a systems enable flexible timing of the onset of gene perturbation in cells already expressing sgRNAs. This feature is particularly important for experiments in microglia: it enables lentiviral delivery of sgRNAs to occur in iPSCs, which are much more amenable to lentiviral infection than microglia, without prematurely affecting genes that may be relevant for differentiation. In the constitutive CRISPRi line, the expression cassette contains a CAG promotor-driven dCas9-BFP-KRAB. In the inducible CRISPRi cassette, this CRISPRi machinery is flanked on both the N and the C termini with dihydrofolate reductase (DHFR) degrons. In the absence of the small molecule trimethoprim (TMP), DHFR degrons cause proteasomal degradation of fused proteins. Addition of TMP stabilizes the degron-tagged CRISPRi machinery. The inducible CRISPRa machinery consists of a DHFR-dCas9-VPH construct, which is similarly stabilized in the presence of TMP.

**Figure 3:**
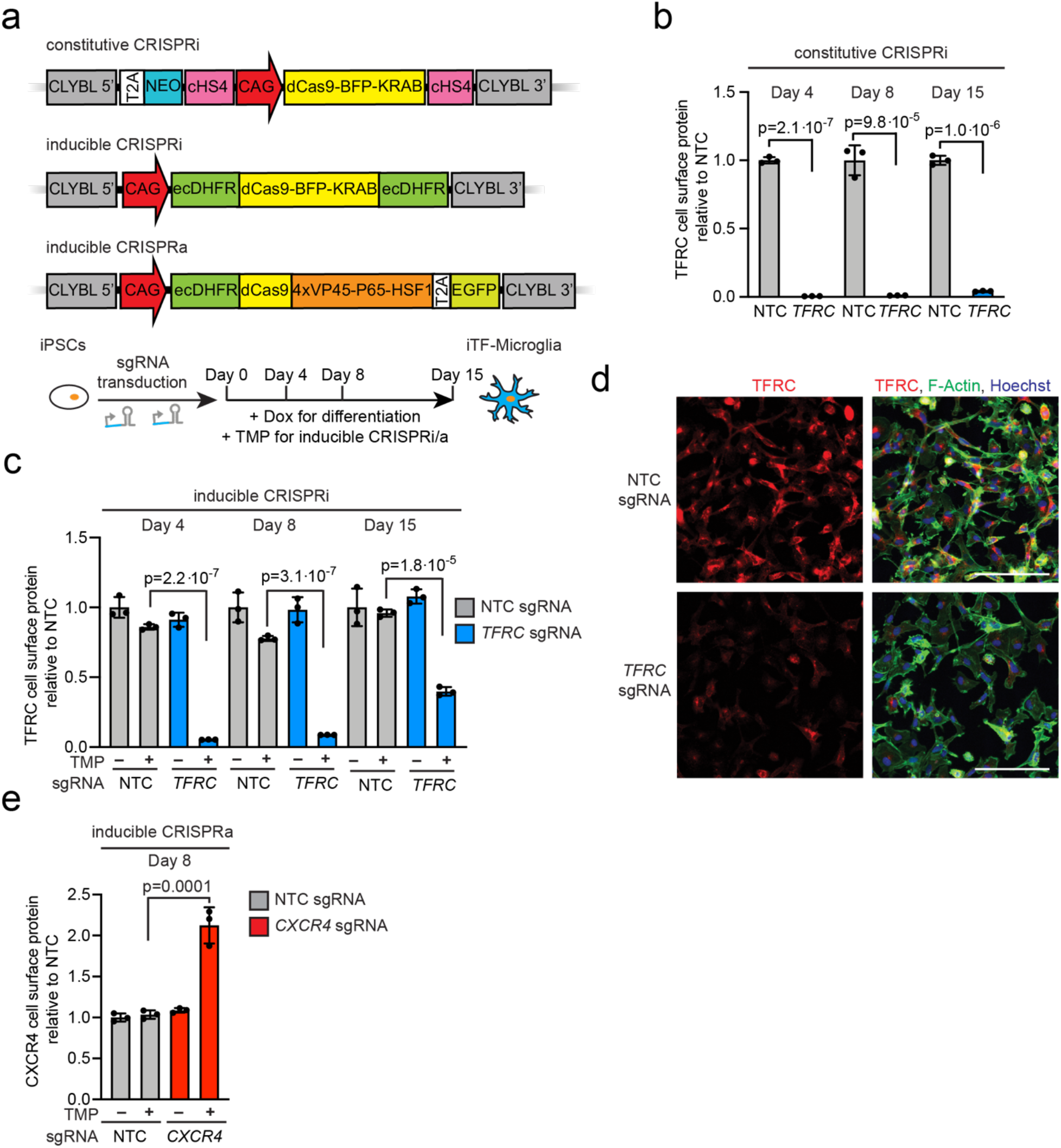
Gene knockdown and overexpression by CRISPRi and CRISPRa in iTF-Microglia. **a,** Strategies for constitutive and inducible CRISPR interference (CRISPRi)/CRISPR activation (CRISPRa) in iTF-Microglia. *Top*: For constitutive CRISPRi, a dCas9-BFP-KRAB construct (catalytically dead Cas9 (dCas9) fused to BFP and the KRAB transcriptional repressor domain) is expressed from the constitutive CAG promotor integrated into the CLYBL safe-harbor locus. *Middle*: For inducible CRISPRi, dCas9-BFP-KRAB is tagged with ecDHFR degrons. *Bottom*: For inducible CRISPRa, CAG promotor-driven ecDHFR-dCas9-VPH was stably integrated into the CLYBL locus. VPH, activator domains containing 4X repeats of VP48, P65 and HSF1. Addition of trimethoprim (TMP) stabilizes the inducible CRISPRi/a machineries. **b,c,** Functional validation of (b) constitutive or (c) inducible CRISPRi activity via flow cytometry of TFRC surface protein level stained iTF-Microglia expressing a TFRC-targeting sgRNA or a non-targeting control (NTC) sgRNA at different days of differentiation (mean +/- sd, n = 3 biological replicates; p values from two-tailed Student’s t-test). (c) TMP was added to induce CRISPRi activity where indicated. **d,** Functional validation of inducible CRISPRi activity via TFRC immunofluorescence (IF) microscopy on Day 8. Top row, non-targeting (NTC) sgRNA. Bottom row, sgRNA targeting *TFRC*. TFRC: red, F-actin: green, nuclei: blue. Scale bar = 100 μm. **e,** Functional validation of inducible CRISPRa activity via flow cytometry of CXCR4 surface protein level staining in iTF-Microglia expressing *CXCR4* sgRNA or non-targeting control (NTC) sgRNA (mean +/- sd, n = 3 biological replicates; p values from two-tailed Student’s t-test). TMP was added to induce CRISPRa activity where indicated.

To validate CRISPRi activity, we transduced iPSCs with a lentiviral construct expressing a sgRNA targeting the transferrin receptor gene (*TFRC*) or a non-targeting control (NTC) sgRNA. In cells expressing the constitutive CRISPRi machinery, knockdown of *TFRC* was robust in iPSCs and iTF-Microglia both on the protein level (Fig. 3b, Extended Data Fig. 4a) and mRNA level (Extended Data Fig. 4c,e). In cells expressing the inducible CRISPRi machinery, TFRC knockdown was completely dependent on the presence of TMP, and effective on the mRNA and protein levels, albeit with reduced knockdown compared to the constitutive CRISPRi system (Fig. 3c,d, Extended Data Fig. 4 b,d,f). For additional target genes we tested, we found examples of excellent around 80% knockdown with both the constitutive and the inducible system for *INPP5D* (Extended Data Fig. 4g,h), but also an example of a gene (*PICALM*) that was effectively knocked down by 90% with the constitutive CRISPRi (Extended Data Fig. 4i), but not by inducible CRISPRi (Extended Data Fig. 4j). Despite these limitations of our current inducible CRISPRi system, we decided to use it for the studies presented in this paper, since it enabled us to induce CRISPRi knockdown only upon differentiation, rather than in the iPSC state, thus reducing the likelihood of recovering phenotypes due to effects in iPSCs or on differentiation itself.

Next, we validated the functionality of the inducible CRISPRa machinery by testing the induction of the endogenous gene *CXCR4*. We observed a robust and tightly inducible increase of *CXCR4* levels in iPSCs and iTF-Microglia on the mRNA level (Extended Data Fig. 4l,m) and the protein level (Fig. 3e, Extended Data Fig. 4k).

### Identification of modifiers of microglial survival/proliferation by CRISPRi screens

Our first application of the inducible CRISPRi iTF-Microglia platform was to identify modifiers of microglia survival and proliferation in a pooled genetic screen (Fig. 4a). First, we transduced the iPSCs with our next-generation lentiviral CRISPRi sgRNA library targeting the “druggable genome”^33^. This library consists of sgRNAs targeting 2,325 genes encoding kinases, phosphatases, and other classes of druggable proteins with five sgRNAs per gene and 500 non-targeting control sgRNAs. After library transduction, the iPSCs were differentiated into iTF-Microglia by addition of doxycyline and TMP was added to induce CRISPRi activity. iTF-Microglia were collected before differentiation (Day 0) and on Day 15 post-induction. Frequencies of cells expressing each sgRNA were determined by next-generation sequencing (Fig. 4a, Supplementary Table 3).

**Figure 4:**
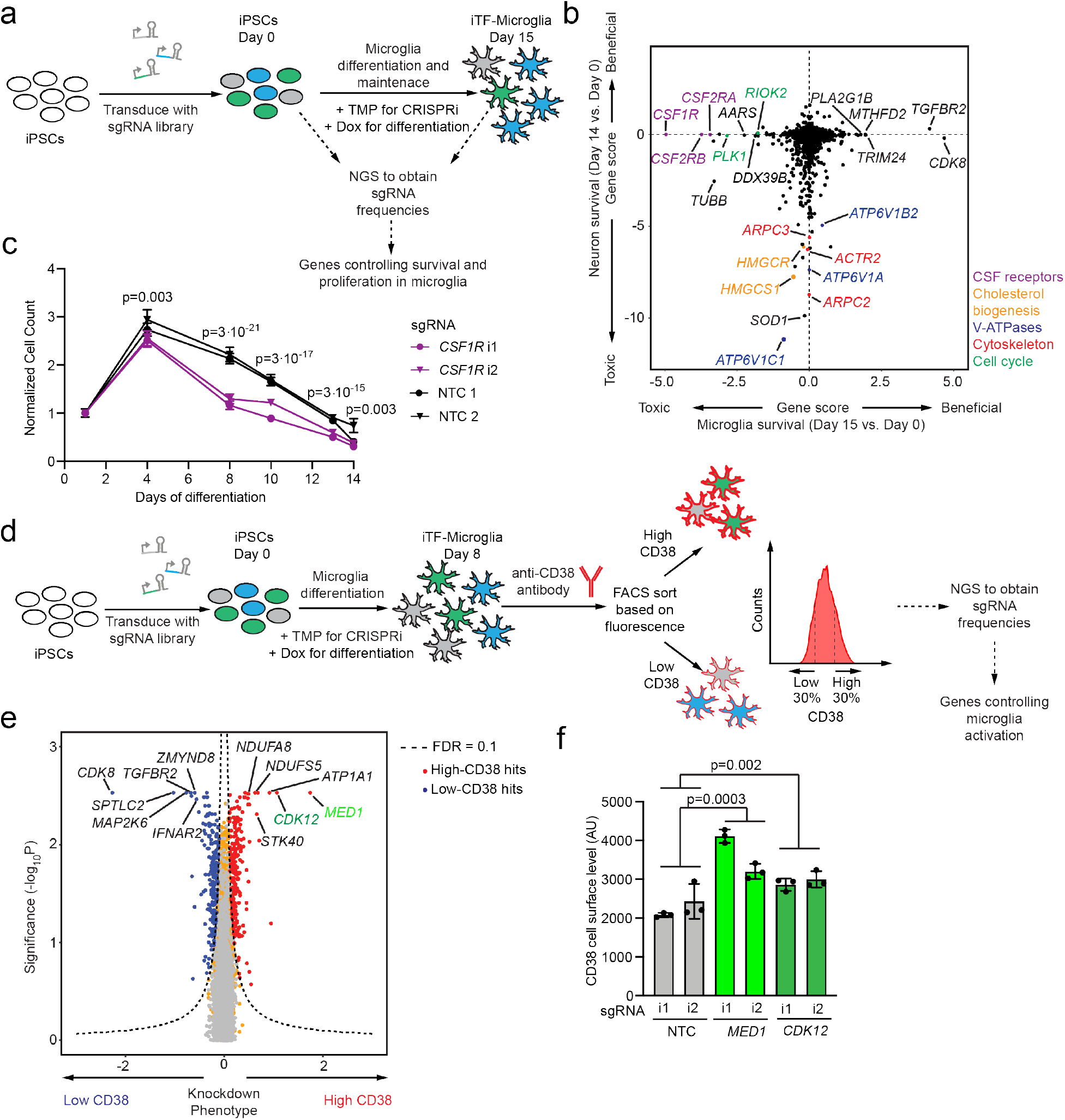
Identification of modifiers of survival and inflammation by CRISPRi screens. **a,** Strategy for the CRISPRi screen to identify modifiers of survival/proliferation. iPSCs expressing the inducible CRISPRi construct were transduced with an sgRNA library targeting the “druggable genome”. On Day 0, doxycyline and cytokines were added to induce microglial differentiation and TMP was added to induce CRISPRi activity. Samples of cell populations were taken at Day 0 and at Day 15 and frequencies of the cells expressing a given sgRNA were determined by next-generation sequencing (NGS) to calculate Gene Scores quantifying the survival/proliferation phenotype for each gene knockdown. **b,** Comparison of Gene Scores from CRISPRi survival screens in iTF-Microglia (this study) vs. iPSC-derived neurons^15^. Each dot represents a gene; genes are color-coded by pathways. **c,** Validation of the phenotype of CSF1R knockdown. iTF-Microglia transduced with *CSF1R*-targeting or non-targeting control (NTC) sgRNAs were imaged on different days after differentiation, and live cells were quantified based on staining with Hoechst 33342. Data is shown as mean +/- sd, n = three wells per group, 7 fields were imaged for each well. **d,** Strategy for a CRISPRi screen to identify modifiers of the expression of CD38, a marker of reactive microglia. iPSCs expressing the inducible CRISPRi construct were transduced with the druggable genome sgRNA library. On Day 0, doxycyline and cytokines were added to induce microglial differentiation, and TMP was added to induce CRISPRi activity. On Day 8, iTF-Microglia were stained for cell-surface levels of CD38 and sorted by FACS into populations with low (bottom 30%) and high (top 30%) CD38 levels. Frequencies of iTF-Microglia expressing a given sgRNA were determined in each population by NGS. **e,** Volcano plot indicating knockdown phenotype and statistical significance (Mann-Whitney U test) for genes targeted in the CD38 level screen. Dashed line indicates the cut-off for hit genes (FDR = 0.1). Hit genes are shown in blue (knockdown decreases CD38 level) or red (knockdown increases CD38 level), non-hit genes are shown in orange and “quasi-genes” generated from random samples of non-targeting control sgRNAs are shown in grey. Hits of interest are labeled. **f,** Validation of the phenotype of *MED1* and *CDK12* knockdown. CD38 cell surface levels measured by flow cytometry of Day 8 iTF-Microglia targeting *MED1*, *CDK12* compared to NTC sgRNA. n = 3 biological replicates; p values from two-tailed Student’s t-test.

We compared the results from the iTF-Microglia survival screen to our previously published^15^ CRISPRi survival screen in iPSC-derived neurons (Fig. 4b) and iPSCs (Extended Fig. 5a). We found that genes affecting microglial survival, neuronal survival and iPSC survival were largely distinct. Knockdown of cholesterol biogenesis enzymes and V-ATPase subunits drastically reduced neuronal but not microglial survival (Fig. 4b). Conversely, knockdown of members of the colony stimulating factor (CSF) receptor family (*CSF1R*, *CSF2RB*, *CSF2RA*) strongly reduced survival of microglia but not neurons (Fig. 4b) or iPSCs (Extended Fig. 5a), consistent with their role in the development and survival of microglia and macrophages^34–37^. We validated CSF1R essentiality in a time-course experiment (Fig. 4c). The toxicity of CSF1R knockdown became pronounced only in differentiated iTF-Microglia (from Day 8 onwards), consistent with the microglia-specific role of CSF1R.

Interestingly, the knockdown of several genes, including *CDK8* and *TGFBR2,* increased abundance of iTF-Microglia in our screen (Fig. 4b). However, we found that *CDK8* and *TGFBR2* knockdown resulted in decreased levels of microglia marker IBA1 (Extended Data Fig. 5b,c), suggesting disrupted microglial differentiation. Indeed, inhibition of TGF-*β* signaling has been shown to compensate for loss of *Oct4* pluripotency signaling^38^ and microglia have been shown to be absent in TGF-*β*1-deficient mice^39^. *CDK8* expression has been shown to correlate to the stem cell pluripotency state and loss of CDK8 could cause iPSCs to differentiate into a non-microglia state^40^. This disruption of the microglia differentiation does not seem specific to our iTF-Microglia differentiation protocol, since knockdown of *CDK8* also decreased IBA1 levels in iPSC-derived microglia we generated using a non-transcription factor-based differentiation protocol^22^ (Extended Data Fig. 5d,e).

To test whether *CDK8* and *TGFBR2* knockdown would also act in differentiated microglia, in addition to their effect on differentiation, we induced their CRISPRi knockdown on Day 8 (Extended Data Fig. 5f). Induction of *CDK8* and *TGFBR2* knockdown in fully differentiated iTF-Microglia did not result in proliferation, and in the case of *TGFBR2* knockdown even in a very slight decrease in survival. By contrast, knockdown of *CSF1R* in Day 8 iTF-Microglia reproduced the phenotype observed in the initial screen (Extended Data Fig. 5f).

### Identification of modifiers of microglial activation by CRISPRi screens

In a second screen, we aimed to identify modifiers of inflammatory activation of microglia. For this screen, we chose cell surface levels of CD38 as a readout for microglial activation. CD38, also known as cyclic ADP ribose hydrolase, is a plasma membrane glycoprotein of all brain cells^41^. CD38 expression and its enzymatic activity increases after LPS and interferon-gamma treatment in primary microglia^42^. Likewise, we observed transcript-level upregulation of CD38 in response to LPS treatment in iTF-Microglia and microglia we differentiated based on a different protocol^22^ (Extended Data Fig. 2e). Similarly, cell-surface levels of CD38 protein increased upon LPS treatment based on flow cytometry (Extended Data Fig. 5g). CD38 plays several roles in microglial activation, including in the secretion of proinflammatory cytokines^43^ and in activation-mediated cell death^42^. Altogether, these data suggest that CD38 is both a marker and an important effector for the activation of microglia and is therefore a suitable marker for a screen for inflammation modifiers.

The screen for modifiers of microglial activation was conducted as shown in Figure 4d. Briefly, iPSCs expressing the inducible CRISPRi machinery were transduced with the pooled sgRNA library described above. The cells were then differentiated into iTF-Microglia, stained for cell surface CD38 using a fluorescently tagged antibody and subjected to FACS sorting into CD38^low^ and CD38^high^ populations. Frequencies of cells expressing each sgRNA were identified in these populations using next-generation sequencing (Supplementary Table 3).

This CRISPRi screen identified several genes regulating cell surface levels of CD38. Knockdown of two transcriptional regulators, *CDK12* and *MED1*, significantly increased CD38 surface levels in the screen (Fig. 4e) and in validation experiments (Fig. 4f). CDK12 is known to be involved not only in cell cycle progression but also in TNF^44^ and noncanonical NF-κB^45^ signaling. While these previous reports may suggest a pro-inflammatory role of CDK12, our findings suggest that the role of CDK12 may be more nuanced or context-dependent, and we designated it for further investigation (see below). Another class of hits whose knockdown increased CD38 levels were members of the mitochondrial Complex I (NADH:ubiquinone oxidoreductase) NDUFA8 and NDUFS5 (Fig. 4e). Knockdown of components of this complex have previously been shown to promote an inflammatory state in macrophages^46^, validating our findings.

Taken together, our large-scale CRISPRi screens in iTF-Microglia uncovered microglia-specific survival modifiers and novel modulators of inflammatory activation, demonstrating the ability of the iTF-Microglia screening platform to identify microglia-specific biology.

### Modifiers of synaptosome phagocytosis by microglia

Microglial phagocytosis is central to brain homeostasis from development through aging^47^. Dysfunction in efferocytosis, the phagocytosis of dead cells, debris, protein aggregates and in synaptic pruning, the phagocytic elimination of neuronal synapses by microglia, have been implicated in neurodegenerative and psychiatric diseases^48–51^. To uncover regulators of microglial phagocytosis, we conducted parallel CRISPRi and CRISPRa screens in iTF-Microglia transduced with sgRNA libraries targeting the “druggable genome”. After 1.5 hours of incubation with pHrodo Red-labeled synaptosomes isolated from rat brains, iTF-Microglia were sorted via FACS based on the pHrodo Red fluorescence signal (Fig. 5a), and screens were analyzed as described for the CD38 FACS-based screen.

**Figure 5:**
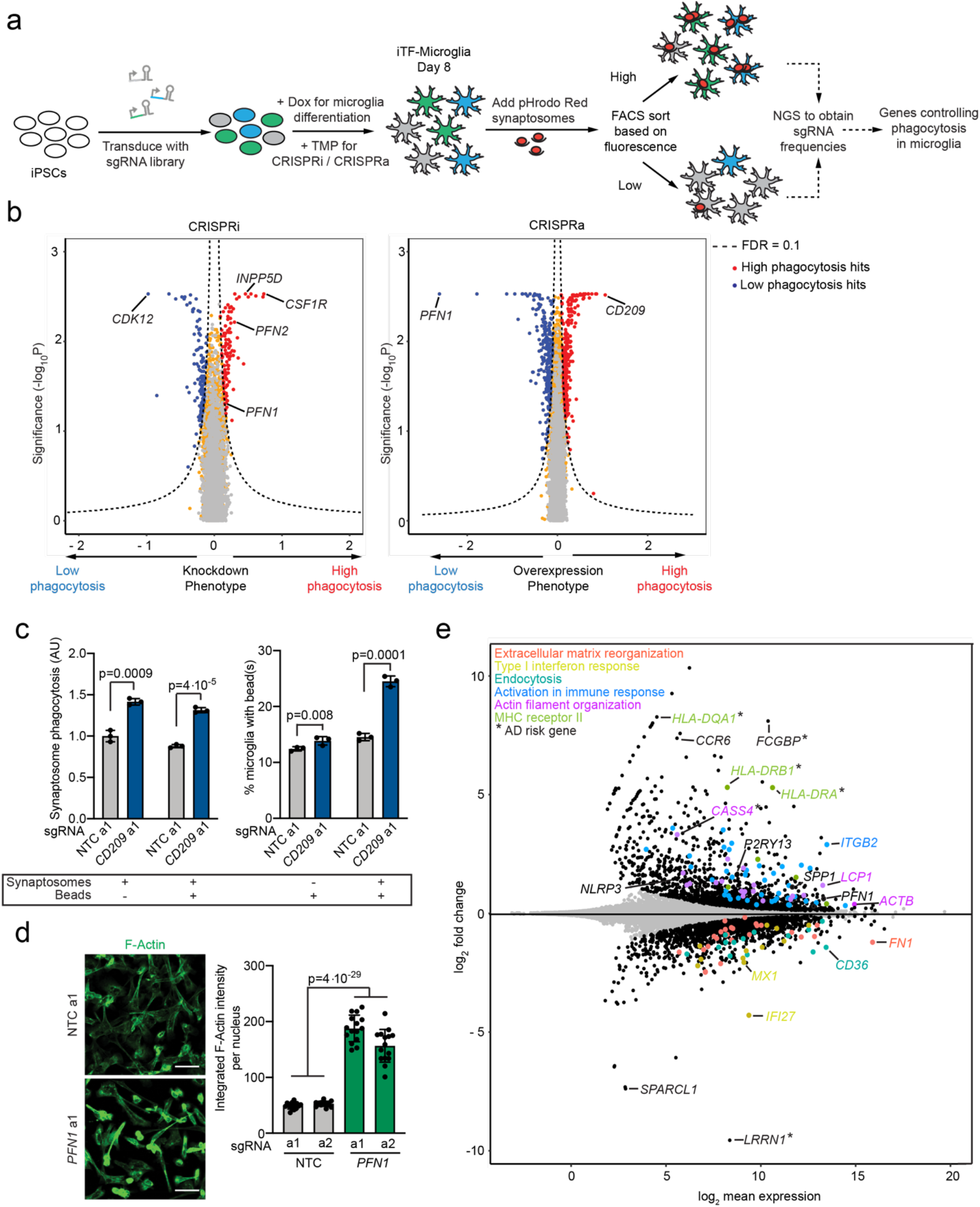
Identification of modifiers of phagocytosis by CRISPRi and CRISPRa screens. **a,** Schematic of the screening strategy to identify modifiers of synaptosome phagocytosis. iPSCs expressing inducible CRISPRi or CRISPRa constructs were transduced with an sgRNA library targeting the druggable genome. On Day 0, doxycyline and cytokines were added to induce microglial differentiation and TMP was added to induce CRISPRi activity. On Day 8, rat synaptosomes labeled with pHrodo Red were added to the cells for 1.5 h and iTF-Microglia were sorted based on fluorescence. Frequencies of cells expressing a given sgRNA in the low-fluorescence and high-fluorescence populations were determined by next-generation sequencing (NGS). **b**, Volcano blots summarizing knockdown and overexpression phenotypes and statistical significance (Mann-Whitney U test) for genes targeted in the pooled phagocytosis screens. Left, CRISPRi screen: right, CRISPRa screen. Dashed lines: Gene Score cutoff for hit genes (FDR = 0.1). Hit genes are shown in blue (knockdown decreases phagocytosis) or red (knockdown increases phagocytosis), non-hit genes are shown in orange and “quasi-genes” generated from random samples of non-targeting control sgRNAs are shown in grey. Hits of interest are labeled. **c,** Competitive phagocytosis assay to test substrate specificity of *CD209* overexpression. Flow cytometry measurement of phagocytosis of pHrodo-Red-labelled synaptosomes (*Left*, either synaptosomes alone or together with beads) and green, fluorescent beads (*Right*, either beads alone or together with synaptosomes) by iTF-Microglia expressing either non-targeting control (NTC) sgRNAs or sgRNAs targeting *CD209*. Values represent mean +/- sd of n=3. Data was analyzed using two-tailed Student’s t-test. **d**, Representative fluorescent images demonstrating higher F-actin staining in CRISPRa iTF-microglia at Day 8 with *PFN1* sgRNAs compared to non-targeting control (NTC) sgRNAs (left). Right, integrated F-actin intensity per cell of CRISPRa iTF-Microglia at Day 8 with *PFN1* sgRNAs or non-targeting control (NTC) sgRNAs. (mean +/- sd, n = 5 fields of view from three different wells per sgRNA. P values from two-tailed Student’s t-test. **e**, Transcriptomic changes caused by *PFN1* overexpression in Day 8 iTF-Microglia (n = 3 biological replicates). Differentially expressed genes (padj < 0.05) are labelled in black. Other colors label genes associated with specific pathways that are discussed in the main text. Alzheimer’s disease (AD) risk genes are labelled with an asterisk.

There was little overlap between CRISPRi and CRISPRa hits (Extended Data Fig. 6a, Supplementary Table 3), confirming our previous findings from screens in diverse biological contexts that overexpression and knockdown screens can provide complementary insights^16, 52, 53^. The underlying reasons include (i) that unlike overexpression screens, knockdown screens can only yield phenotypes for genes expressed in the cell type under investigation, and (ii) that knockdown of a single element of a pathway or multi-subunit complex can cause a loss-of-function phenotype, whereas overexpression of a single element is generally not sufficient to elicit a gain-of-function phenotype for an entire pathway or multi-subunit complex. A prominent exception was the actin-binding protein *PFN1*, coding mutations in which cause ALS^54^. *PFN1* had opposing phenotypes on synaptosome phagocytosis upon CRISPRi repression and CRISPRa induction (Fig. 5b). Unexpectedly, knockdown of *CSF1R* increased phagocytosis (Fig. 5b), even though, as we had previously found (Fig. 4b) its knockdown decreased iTF-Microglia survival. Another remarkable hit was the Alzheimer’s disease risk factor *INPP5D*, knockdown of which slightly increased phagocytosis. Overexpression of *CD209*, a C-type lectin receptor present on the surface of macrophages and dendritic cells, greatly increased synaptosome phagocytosis (Fig. 5b). We validated these phenotypes from the primary CRISPRi screen individually in iTF-Microglia (Extended Data Fig. 6b) and in iPSC-derived microglia generated an alternative protocol^22^ (Extended Data Fig. 6c).

We further investigated the CRISPRa hits *PFN1* and *CD209*. We validated upregulation of both genes by qPCR (Extended Data Fig. 6e). Pattern-recognition receptor CD209 has previously been shown to regulate phagocytic capacity in macrophages^55^. We wondered if *CD209* was additionally a substrate-specific phagocytosis regulator. To directly test the effect of *CD209* on substrate specificity, we adapted our phagocytosis assay to simultaneously test uptake of two separate substrates, pHRodo-Red labeled synaptosomes and yellow-green (YG) fluorescently labeled beads (Fig. 5c). Using this approach, we challenged iTF-Microglia with either synaptosomes or beads alone, or a mixture of both. Consistent with the screen result, overexpression of *CD209* increased phagocytosis of synaptosomes. Interestingly, it only changed bead phagocytosis by 10% (Fig. 5c), suggesting substrate specificity. However, when challenging iTF-Microglia with a mixture of beads and synaptosomes, bead phagocytosis was 2-fold increased compared to control iTF-Microglia. This finding suggests that presence of synaptosomes might stimulate general phagocytosis via *CD209*.

In addition to decreased synaptosome phagocytosis, we observed increased F-actin levels in iTF-Microglia as a consequence of *PFN1* overexpression induced by CRISPRa (Fig. 5d), consistent with previous finding that moderate overexpression of PFN1 induces long stress fiber-like actin cables^56^. This process could disturb orchestrated actin polymerization at the membrane and thus decrease phagocytosis. In addition to direct effects on the actin cytoskeleton, *PFN1* knockdown has also been reported to result in anti-inflammatory changes^57^. Indeed, we observed transcriptional changes in immune-related genes and AD risk genes upon *PFN1* overexpression in iTF-Microglia (Fig. 5e, Supplementary Table 4).

In conclusion, our complementary CRISPRi and CRISPRa screens identify known as well as novel phagocytosis modulators in microglia, which validated in iPSC-derived microglia generated using an alternative protocol.

### Single-cell transcriptomics reveal distinct states of iTF-Microglia

Several genes had CRISPRi phenotypes in more than one of the large-scale screens that we conducted (Extended Data Fig. 6f, Supplementary Table 3). We therefore hypothesized that some hit genes were not dedicated factors required for specific microglial processes, but rather more global regulators of distinct functional states. To test this hypothesis and gain more detailed insights into the mechanisms by which genes affect microglial functions, we selected 39 hit genes of interest, most of which had phenotypes in more than one of the large-scale primary screens (Extended Data Fig. 6f, Supplementary Table 3) for characterization in a CROP-seq screen, which couples CRISPRi perturbation to single-cell RNA sequencing. We introduced a library of 81 sgRNAs (two sgRNAs targeting each selected gene (only a single sgRNA was included for *DBF4* due to an error) and four non-targeting control sgRNAs; Supplementary Table 5) into iPSCs, induced iTF-Microglia differentiation and CRISPRi activity, and performed single-cell RNA sequencing of 58,302 iTF-Microglia on Day 8 (Fig. 6a, Supplementary Table 6).

**Figure 6:**
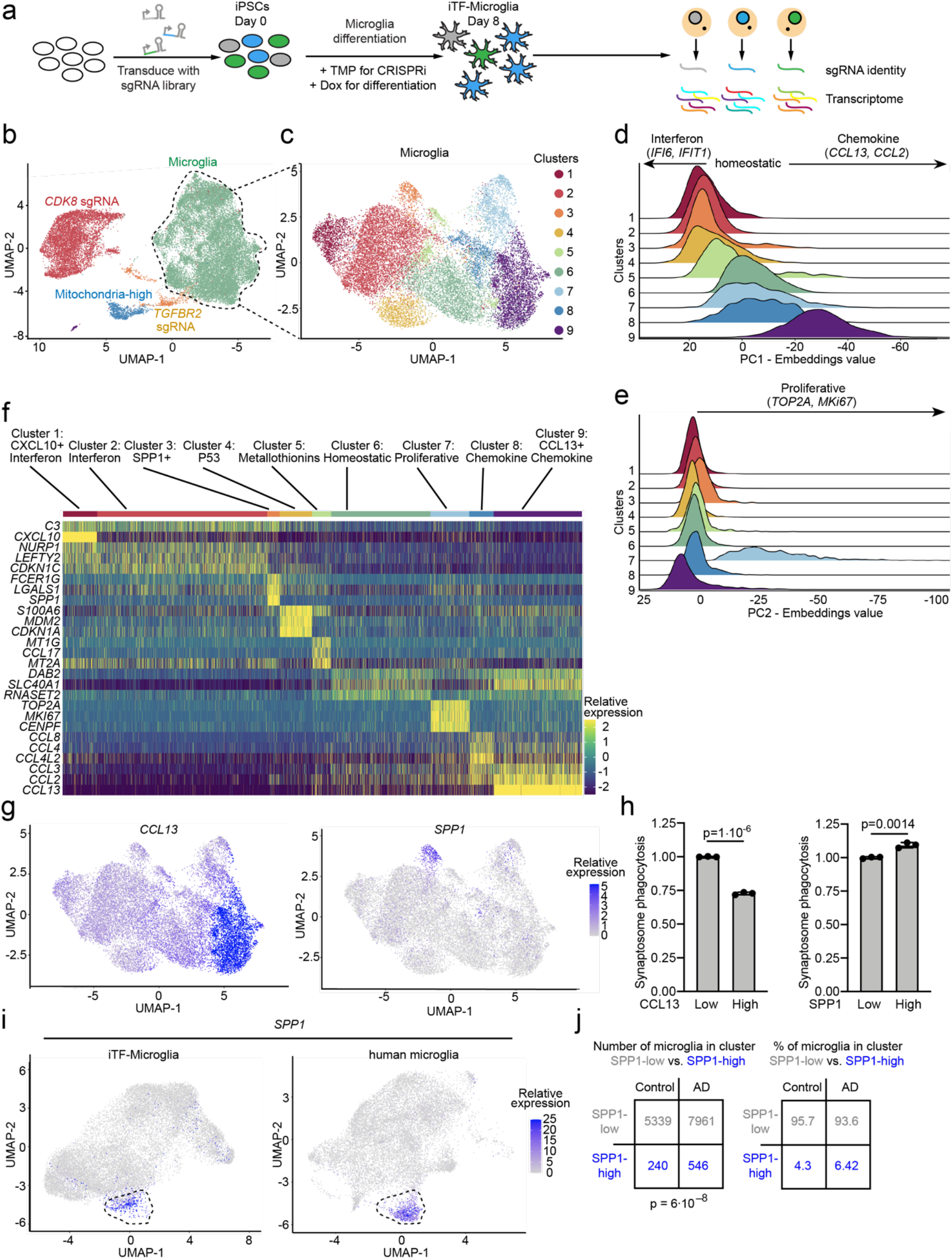
Single-cell RNA sequencing reveals distinct and disease-related microglia subclusters. **a,** Strategy for the CROP-seq screen. IPSCs expressing inducible CRISPRi machinery were transduced with a pooled library of 81 sgRNAs CROP-seq vector pMK1334. iPSCs are differentiated to iTF-Microglia and subjected to scRNAseq to obtain single-cell transcriptomes and identify expressed sgRNAs. **b**, UMAP projection of the 28,905 cells in the post-QC CROP-seq dataset. Cells are colored by sgRNA (*CDK8*-red, *TGFBR2*-orange) and cells with a high percentage of mitochondrial transcripts (blue). Microglia are labeled in green. Each dot represents a cell. **c**, UMAP projection depicting the 9 different clusters within the 19,834 microglia. Each dot represents a cell. The cells are color-coded based on their cluster membership. **d-e**, Ridge plots depicting iTF-Microglia clusters along PC1 (d) and PC2 (e). PC1 spans inflammation status (Interferon activated-homeostatic-chemokine activated) while PC2 spans proliferation status. **f**, Heatmap of iTF-Microglia clusters 1-9 and the relative expression of the top three differentially expressed genes of each cluster. **g**, UMAP projection of distinct marker expression of *CCL13* (*Left*) and *SPP1* (*Right*). *CCL13* is a marker for cluster 9 and *SPP1* is a marker for cluster 3. Cells are colored by the expression levels of the indicated gene. **h**, Phagocytic activity of iTF-Microglia in different states. Flow cytometry measurement of phagocytosis of pHrodo-Red-labelled synaptosomes (*Left*, Phagocytosis in CCL13^high^ and CCL13^low^ iTF-Microglia) (*Right*, Phagocytosis in SPP1^high^ and SPP1^low^ iTF-Microglia). Values represent mean +/- sd of n=3; p values from two-tailed Student’s t-test. **i**, Integration of single-cell transcriptomes of iTF-Microglia and microglia from post-mortem human brains^7^. In the integrated UMAP, iTF-Microglia (*Left)* with high *SPP1* expression and human brain-derived microglia with high *SPP1* expression (*Right*) form a cluster (dashed outline). **j**, In brains from patients with Alzheimer’s disease (AD), a higher fraction of microglia is in the SPP1^high^ cluster compared to control brains (data from Olah *et al*.^7^; P value from two-sided Fisher’s exact test).

Unsupervised clustering and Uniform Manifold Approximation and Projection (UMAP) dimensional reduction of the single-cell transcriptomes uncovered distinct clusters (Fig. 6b). In one cluster, a high proportion of transcripts mapped to mitochondrial transcripts, suggesting damaged or dying cells; this cluster was removed from downstream analysis (Extended Data Fig. 7a). Two clusters exclusively contained cells expressing sgRNAs targeting *CDK8* or *TGFBR2* (Fig. 6b and Extended Data Fig. 7b). These cells expressed high levels of the pluripotency marker *SOX2*, but low levels of the microglia marker *CSF1R* (Extended Data Fig. 7c). Together with our previous experiments showing reduced IBA1 levels for iTF-Microglia targeting *CDK8* and *TGFBR2* (Extended Data Fig. 5b,c), these findings suggest that microglia differentiation was disrupted in those cells. We removed those clusters from further analysis and retained the remaining cluster, which was characterized by high levels of *CSF1R* expression (Extended Data Fig. 7c). Importantly, 92.4% of cells expressing NTC sgRNAs were part of the cluster with high levels of *CSF1R* expression, confirming the high efficiency of our microglial differentiation protocol for unperturbed cells.

Unsupervised clustering and UMAP dimensional reduction of the remaining 19,834 iTF-Microglia revealed 9 transcriptionally distinct clusters (Fig. 6c, Extended Data Fig. 7d). Microglia heterogeneity in response to different environmental conditions in the brain has been extensively studied^58^, but we were surprised to observe a wealth of distinct transcriptional states in the cultured iTF-Microglia. Importantly, NTC sgRNAs are represented in cells in every cluster, suggesting the observed heterogeneity is an innate quality of the iTF-Microglia (Extended Data Fig. 7d).

Principal component analysis identified two major biological axes broadly defining these states. The first principal component (PC-1) corresponded to a polarized axis of inflammatory activation: starting from a central homeostatic state (cluster 6), one direction was defined by interferon-induced gene expression, whereas the other direction was defined by induction of chemokines (Fig. 6d and Extended Data Fig. 7e). The second principal component (PC-2) captures markers of proliferation, mainly in cluster 7 (Fig. 6e and Extended Data Fig. 7f).

To further interpret each transcriptional microglia state, we performed differential gene expression analysis across the clusters (Supplementary Table 7) and named each cluster according to characteristic transcriptomic signatures. Figure 6f highlights the top three genes selectively expressed by cells in each cluster. Clusters 1 and 2 are both defined by high expression of interferon-induced genes and the complement gene *C3.* Cluster 1 is uniquely defined by high expression of chemokine *CXCL10.* Subsets of microglia characterized by upregulation of interferon response genes have been described in mouse models of neurodegeneration^8^.

Cluster 3 is defined by the high expression of *SPP1* (Fig. 6g), which encodes secreted phosphoprotein 1, also known as osteopontin, a multifunctional protein acting both as part of the extracellular matrix and as a secreted cytokine^59^. Importantly, SPP1 is upregulated in several disease-associated microglia states in both human patients and mouse disease models. These include disease-associated microglia (DAM)^9^ and activated response microglia (ARM)^60^ in AD mouse models, and late-response microglia in CK-p25 mouse models of neurodegeneration^8^. SPP1-positive microglia states are also enriched in multiple sclerosis (MS) patients and mouse models^12^ and enriched in microglia in the aging human brain^13^. Furthermore, SPP1 is highly expressed in glioma-associated microglia in mice and humans, where high expression of SPP1 is associated with poor prognosis^61^. Using flow cytometry, we found that SPP1-positive microglia have a slightly increased phagocytic activity, whereas CCL13-positive microglia have substantially decreased phagocytic activity (Fig. 6h). Integration of our iTF-Microglia dataset with a recent scRNA-seq dataset containing 16,242 human microglia from control and Alzheimer’s Disease patient brains^7^ showed conservation of the SPP1-positive microglia state (Fig. 6i). The proportion of SPP1-positive microglia was substantially increased in Alzheimer’s disease patients compared to controls (Fig. 6j). Notably, it remains to be determined how the SPP1+ microglia state affects the pathogenesis of different diseases, since SPP1 has been linked to both pro-inflammatory and anti-inflammatory responses^59^. This question has been challenging to address since we have lacked tools to manipulate the SPP1+ state of microglia *in vivo*.

Cluster 4 is defined by expression of pro-apoptotic p53 signaling genes, and Cluster 5 by expression of metallothionines. Cluster 6 is defined by the absence of interferon response genes or chemokines, and thus we interpreted it as representing homeostatic microglia. Cluster 7 is characterized by the expression of proliferation markers such as *TOP2A* and *MKI67*. Cluster 8 and 9 are characterized by the expression of high levels of chemokines such as *CCL2* and *CCL3*. Cluster 9 is uniquely defined by high expression of *CCL13* (Fig. 6g). Such chemokine signatures have recently been found to be a hallmark of human microglia not observed in mice^62^.

Taken together, single-cell RNA sequencing revealed that many important features of microglia diversity observed in human brains and in disease states are recapitulated in our iTF-Microglia *in vitro* model.

### CROP-seq uncovers regulators of microglial cell states

We next identified the differentially expressed genes (DEGs) caused by CRISPRi knockdown of each gene targeted in the CROP-Seq screen (Extended Data Fig. 8, Supplementary Table 8). As expected, knockdown of functionally related genes shared common DEG signatures. For example, knockdown of *CSF1R*, *CSF2RA* and *CSF2RB* resulted in an upregulation of genes encoding the major histocompatibility complex as well as *CD36*, *CD74* and *CD68* (Extended Data Fig. 8, Supplementary Table 8), which are markers of phagocytic microglia and could possibly explain the increased phagocytic capacity we observed in response to *CSF1R* knockdown (Fig. 5b, Extended Data Fig. 6b).

Given the surprising heterogeneity of iTF-Microglia, we investigated if CRISPRi knockdown of specific genes could control microglial cell states. Indeed, cells containing sgRNAs targeting genes such as *CSF1R*, *CDK12* and *MAPK14* were enriched or depleted from specific clusters (Fig. 7A), and more generally, knockdown of many genes specifically affected the frequency of cell states (Fig. 7b, Extended Data Fig. 9a, Supplementary Table 9).

**Figure 7:**
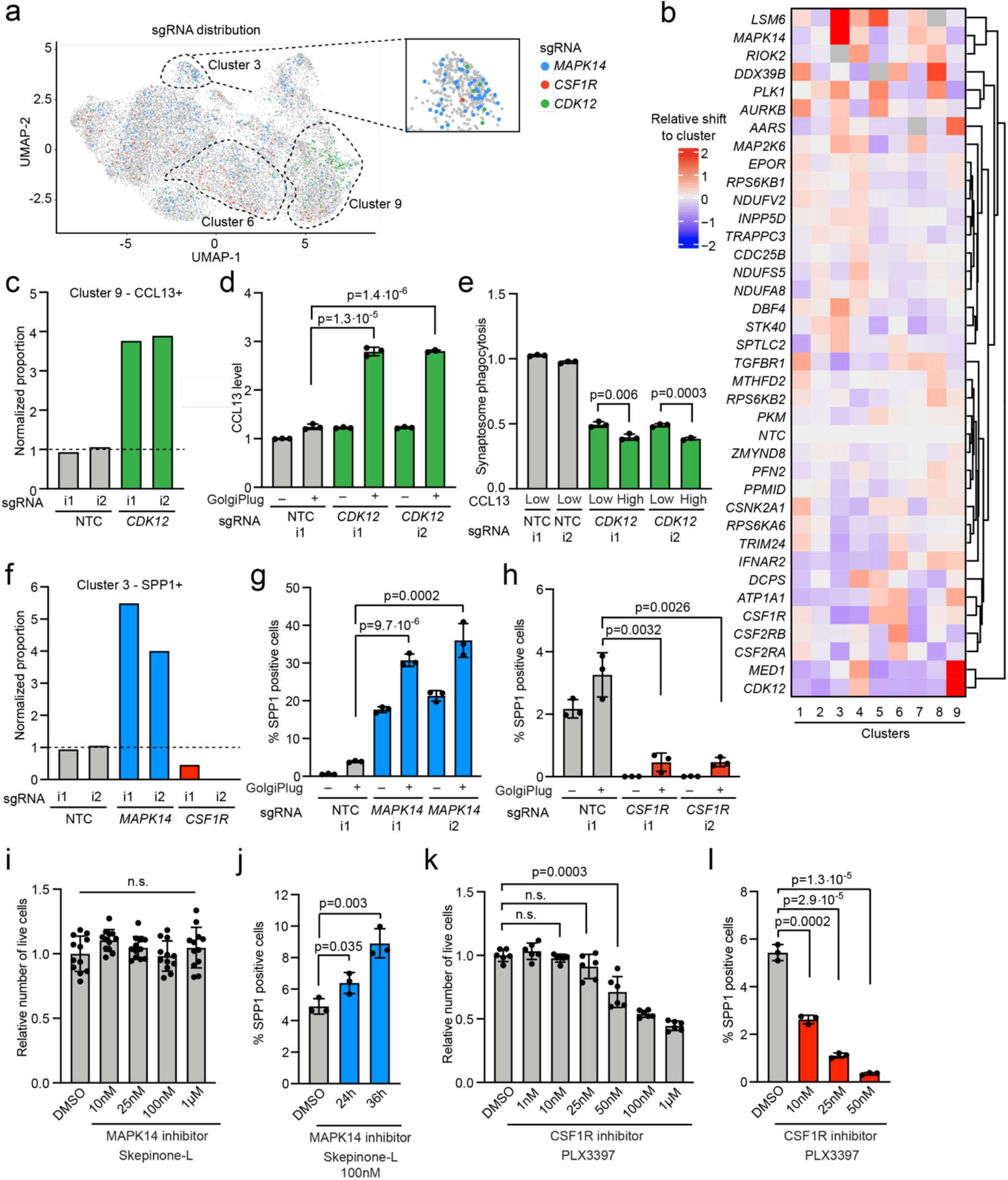
CROP-seq reveals changes in cluster occupancy induced by gene knockdown. **a,** sgRNA distribution across the iTF-Microglia clusters. UMAP projection depicts cells colored by sgRNA. Cells with sgRNAs targeting *MAPK14* (blue), *CSF1R* (red), and *CDK12* (green), are enriched in clusters 3, 6, and 9, respectively. Insert shows cluster 3. **b**, Changes in cluster distribution after CRISPRi knockdown of targeted genes in iTF-Microglia. Heatmap with hierarchical clustering of 37 target genes and non-targeting control (NTC) and their distribution in clusters 1-9. **c,** Proportion of cells in cluster 9 (CCL13+) expressing either non-targeting control (NTC) sgRNAs or sgRNAs targeting *CDK12*. **d**, Functional validation of increased CCL13 level in iTF-Microglias expressing sgRNAs targeting *CDK12* compared to cells expressing a non-targeting control (NTC) sgRNA. CCL13 levels were measured via flow cytometry +/- 5h of GolgiPlug treatment. Values represent mean +/- sd of n = 3 biological replicates; p values from two-tailed Student’s t-test. **e,** Decreased synaptosome phagocytosis of iTF-Microglia expressing sgRNAs targeting *CDK12* compared to cells expressing non-targeting control (NTC) sgRNA. Phagocytosis is further reduced in the CCL13-high population of cells expressing sgRNAs targeting *CDK12*. Phagocytosis was measured via flow cytometry with additional staining for CCL13. Values represent mean +/- sd of n = 3 biological replicates; p values from two-tailed Student’s t-test. **f,** Proportion of cells in cluster 3 (SPP1+) expressing either non-targeting control (NTC) sgRNAs or sgRNAs targeting *MAPK14* or *CSF1R.* **g-h,** Functional validation of altered percentage of SPP1 positive cells in iTF-Microglias expressing sgRNAs targeting *MAPK14* (g) or CSF1R (h) compared to cells expressing a non-targeting control (*NTC*) sgRNA. SPP1 was measured via flow cytometry after treating cells for 5h with GolgiPlug. Values represent mean +/- sd of n = 3 biological replicates; p values from two-tailed Student’s t-test. **i,** Survival of iTF-Microglia after treatment with various concentrations of MAPK14 inhibitor Skepinone-L. Viable cells were quantified using the CellTiter-Glo assay 24h after treatment with Skepinone-L. Values represent mean +/- sd of n = 12 biological replicates. Data was analyzed by ANOVA. **j**, Percentage of SPP1-positive cells after 100 nM Skepinone-L treatment. SPP1 was measured via flow cytometry after treating cells for 24 h or 36 h with Skepinone-L and an additional 5 h with GolgiPlug. Values represent mean +/- sd of n = 3 biological replicates; p values from two-tailed Student’s t-test. **k**, Survival of iTF-Microglia after treatment with various concentrations of CSF1R inhibitor PLX3397. Viable cells were quantified using the CellTiter-Glo assay, 24h after treatment with PLX3397. Values represent mean +/- sd of n = 6 biological replicates; p values from two-tailed Student’s t-test. **j,** Percentage of SPP1 positive cells after PLX3397 treatment. SPP1 was measured via flow cytometry after treating cells for 24 h with PLX3397 and an additional 5 h with GolgiPlug. Values represent mean +/- sd of n = 3 biological replicates; p values from two-tailed Student’s t-test.

Knockdown of *CDK12* shifted cells into cluster 9 (CCL13+, chemokine) (Fig. 7a-c). To validate this phenotype, we used a flow-cytometry approach in which secretion of CCL13 was inhibited with the transport inhibitor GolgiPlug. CCL13 levels in were increased over 2-fold with knockdown of *CDK12* (Fig. 7d), confirming our screen results. Next, we asked if the shift into cluster 9 might also have functional consequences for the iTF-Microglia. We measured synaptosome phagocytosis in CCL13^high^ cells (representative of the cells in cluster 9) and CCL13^low^ cells (representative of other clusters) in *CDK12* knockdown iTF-Microglia. As observed already in our phagocytosis screen (Fig. 5b), knockdown of *CDK12* decreased synaptosome phagocytosis, both in the CCL13^low^ and CCL13^high^ population (Fig. 7e). *CDK12* knockdown caused transcriptional downregulation of phosphatidylserine recognition receptors in both CCL13^low^ and CCL13^high^ cells, which may contribute to the decreased phagocytic activity (Extended Data Fig. 9b). Phagocytosis was even further decreased in the CCL13^high^ population (Fig. 7e), suggesting that microglia in the CCL13+ state have lower phagocytic capacity, as we showed previously (Fig. 6h). Knockdown of *CDK12* also had some cluster-specific effects (Extended Data Fig. 9b), highlighting the complex effects of gene perturbation in both shifting occupancy of cells between defined functional states, but also affecting cellular pathways in both general and state-specific ways. Interestingly, *MED1* had very similar knockdown phenotypes to *CDK12*, and both genes encode factors associated with general transcription by RNA polymerase II.

We next turned our attention to regulators of the disease-relevant SPP1-positive cluster 3. Knockdown of *MAPK14* and *CSF1R* had dramatically opposing effects, increasing and decreasing occupancy in the SPP1 cluster, respectively (Fig. 7a,b,f). Using GolgiPlug treatment to block secretion of SPP1, we validated these phenotypes by flow cytometry: knockdown of *MAPK14* increased the population of SPP1+ cells more than 6-fold (Fig. 7g), whereas CSF1R knockdown greatly diminished the proportion of SPP1+ cells (Fig. 7h).

Based on these results of genetic perturbations, we asked if pharmacological targeting of the same hits would similarly modulate the abundance of the SPP1+ state. Indeed, inhibition of MAPK14 with Skepinon-L increased the fraction of SPP1+ microglia in a time-dependent manner at nontoxic concentrations (Fig 7i,j)

Given that pharmacological inhibition of CSF1R has shown beneficial effects in several neurodegenerative mouse models, and was observed by us and others to selectively affect subpopulations of microglia in mice^63–65^, we tested if pharmacological inhibition of CSF1R would reduce the proportion of SPP1+ microglia. While the CSF1R inhibitor PLX3397 showed dose-dependent toxicity in iTF-Microglia (Fig. 7k), low concentrations of CSF1R inhibitor that were nontoxic or low toxic to bulk iTF-Microglia selectively depleted SPP1+ iTF-Microglia (Fig. 7l). Thus, both pharmacological and genetic inhibition of CSF1R can decrease the proportion of SPP1+ cells.

In conclusion, our CROP-seq screen enabled deep characterization of the hit genes that our primary screens identified and revealed the existence of a wealth of microglia cell states and their regulators. To enable the scientific community to further explore this large dataset, we implemented additional functionality in the CRISPRbrain data commons (https://www.crisprbrain.org/) we previously described^16^. Specifically, interactive three-dimensional UMAP representations and heatmaps enable the selective investigation of cells by expression levels of genes of interest, sgRNA identity, and cluster membership.

## DISCUSSION

In this study, we described a novel platform for large-scale, multimodal CRISPRi/a-based genetic screens in human iPSC-derived microglia. We demonstrated the power of this platform in multiple large-scale screens. While CRISPR knockout strategies are commonly used for loss-of-function screens, the partial knockdown achieved by CRISPRi enables a more nuanced characterization of the function of essential genes. For example, we uncovered a selective vulnerability of microglia in the SPP1+ state to partial knockdown of the microglia-essential gene *CSF1R*. The use of human microglia (as opposed to mouse primary microglia) enabled us to recapitulate microglia features found in human but not mouse brain, such as a state characterized by a chemokine signature^62^. iPSC technology will also make it possible to conduct CRISPRi/a screens in patient-derived cells to identify modifiers of phenotypes linked to genetic risk variants^14^.

Notwithstanding, there are several areas for future optimization of our iTF-Microglia platform. Improved inducible CRISPRi/a machinery with more potent gene repression and activation in fully differentiated iTF-Microglia would enable the induction of CRISPRi/a at later stages during differentiation to avoid false-positive hits that affect microglial differentiation, such as *CDK8* and *TGFBR2* (Fig. 4b, Extended Data Fig. 5b-f).

Another goal for future technology development is further acceleration and enhancement of the microglial maturation. One potential concern about sustained expression of transgenic transcription factors is that this could promote certain microglial states over others. A protocol in which transcription factor expression is discontinued after day 8 (Extended Data Fig. 1a) can mitigate this concern. As with all currently available *in vitro* culture systems, microglia are slightly activated in monoculture and lose their unique homeostatic brain signature^32^. Previous research has shown that iPSC-microglia become more homeostatic in co-culture with neurons^21^, which is compatible with our own observation of enhanced ramification of iTF-Microglia in neuronal co-culture (Fig. 2f). Alternatively, optimizing the set of transcription factors used to generate iTF-Microglia may result in improved abundance of homeostatic microglia. CRISPRa screens in our current platform are a scalable strategy to identify additional transcription factors to promote microglial maturation and homeostasis, leading to ever more faithful models of human microglia.

While there is room for future improvements, our current platform already uncovered new insights into microglial biology. We identified several genes associated with neurodegenerative diseases, including *PFN1* and *INPP5D*, as modulators of phagocytosis in microglia (Fig. 5), thus pointing to a possible cellular mechanism by which variants in these genes contribute to disease. Coding mutations in profilin 1 (*PFN1*) gene cause amyotrophic lateral sclerosis (ALS)^54^. *PFN1* is a small actin-binding protein that promotes formin-based actin polymerization and regulates numerous cellular functions, but how mutations in *PFN1* cause ALS is unclear. The actin cytoskeleton is known to be important for the physiological functions of microglia, including migration and phagocytosis. We observed that *PFN1* overexpression disrupts the actin cytoskeleton in iTF-Microglia with higher levels of F-actin. Recently, a study has shown that *PFN1* is also involved in microglia activation, since knockdown of *PFN1* inhibited M1 proinflammatory microglial polarization and promoted anti-inflammatory M2 microglia polarization after oxygen and glucose deprivation^57^. Introducing the ALS-associated mutations in the *PFN1* gene in iPSCs will shed light on the impact of these specific mutations on the function of different relevant cell types, such as iPSC-derived neurons and microglia.

Genetic variants in the *INPP5D* locus are associated with an increased susceptibility to AD^66^ and cerebrovascular function as well as tau and A*β* levels in the cerebrospinal fluid of AD patients^67^. *INPP5D* encodes the lipid phosphatase SHIP1, which is selectively expressed in brain microglia. SHIP1 inhibits signal transduction initiated by activation of immune cell surface receptors, such as TREM2^68^. Intriguingly, *INPP5D* expression increases with AD progression, predominantly in plaque-associated microglia, and correlates with plaque density^69^. Given the results from our phagocytosis screen, *INPP5D* overexpression might result in microglia with deficient phagocytic capacity, resulting in increased Aβ deposition and neurodegeneration. Concordant with the findings from our genetic screen, a recent study found that pharmacological SHIP1/2 inhibitors promote microglial phagocytosis *in vitro* and *in vivo*^70^.

Our single-cell RNA sequencing screen revealed that iTF-Microglia adopt a spectrum of states, including states mirroring those observed in human brains, such as the SPP1-positive state. Even though our protocol uses overexpression of transcription factors to generate microglia-like cells, our combination of six transcription factors does not specify a single state, but recapitulates the intrinsic plasticity of cell states that is a hallmark of microglia.

Our CROP-Seq screen identified genes controlling the distribution of iTF-Microglia across distinct cell states. Knockdown or pharmacological inhibition of *MAPK14* strongly promoted adoption of the disease-associated SPP1-positive state. Previous work suggested a functional connection between SPP1 and MAPK14 in cancer cells, where SPP1 can activate the p38 MAPK signaling pathway, which comprises MAPK14^71^. MAPK14 was also recently predicted to be a unique network regulator in DAM^72^. However, our identification of MAPK14 as a regulator of the SPP1+ state is novel and enhances our understanding of modulators of microglia cell states.

We found that the SPP1-positive microglia state can be selectively depleted by genetic and pharmacological inhibition of CSF1R. CSF1R inhibitors have beneficial effects in mouse models of diseases including AD^73, 74^, tauopathy^65^ and MS^75^. Intriguingly, CSF1R inhibition reduced SPP1 expression in the MS model, while homeostatic genes such as *TMEM119* and *P2RY12* were increased^75^, paralleling our finding that the SPP1 microglia state is selectively vulnerable to CSF1R inhibition. Additionally, disruption of CSF1-CSF1R signaling downregulated SPP1 in the cerebellum^76^. Combining CSF1R depletion and single cell profiling has enabled us previously to elucidate the differential effects of CSF1R inhibitors on microglia subtypes^63^. Following CSF1R inhibition, we found an enrichment of microglia states with elevated markers of inflammatory chemokines and proliferation and interestingly, in concordance with our findings in iTF-Microglia here, an upregulation of cell surface receptor CD74^63^. Others have reported compensatory upregulation of TREM2/*β*-catenin and IL-34 in microglia following conditional CSF1R KO^77^; however, we did not find consistent upregulation of these factors in our iTF-microglia (Supplementary Table 9). Based on our new finding that CSF1R inhibition at low doses that are nontoxic to most microglia selectively depletes the SPP1+ population in iTF-Microglia, low-dose CSF1R inhibition might also give us a tool to study the SPP1+ population in mouse disease models.

Taken together, our results have provided pharmacological strategies to either promote or deplete SPP1+ microglia. This will make it possible to determine the role of SPP1+ microglia in different diseases, where they may play either beneficial or detrimental roles, and to manipulate this disease-associated microglial state for therapeutic benefit.

We anticipate that the screening platform we describe here can be broadly applied to screen for other microglia-related phenotypes, and to systematically identify regulators of different microglia states. Using iPSCs derived from patients with familial or sporadic diseases will enable the identification of potential therapeutic targets that can correct cellular phenotypes. For example, the APOE ε4 variant reduces the ability of iPSC-microglia to clear Aβ^78^ and dysregulates cholesterol biogenesis^79^, and CRISPR screens have the potential to elucidate the underlying molecular mechanisms and uncover potential remedies. Introduction of microglia into co-cultures or brain organoids can provide a screening platform to investigate their interactions with other brain cell types, such as synaptic pruning of neurons. Finally, transplantation of iTF-Microglia into postnatal, immune-deficient, humanized mice could result in microglia with an *ex vivo* human microglial gene signature, including more homeostatic microglia, and enable the investigation of factors controlling the interaction of microglia with a model for diseased brain environment^80–83^.

## ACKNOWLEDGEMENTS

We thank Ruilin Tian for discussions about CROP-Seq, Chris Bohlen for advice on synaptosome protocols, Jessica Mella, Karinna Vivanco, Capria Rinaldi, Gabriel Sturm, Yaqiao Li, Carlo Condello, and Miranda Sullivan for contributions to preliminary studies. We thank Avi Samelson for feedback on the manuscript and members of the Kampmann lab for discussions.

This research was supported by NIH grants DP2 GM119139 and U01 MH115747 to MK, U54 NS100717 to LG and MK, R01 AG051390 to LG, F30 AG066418 to KL, F30 AG062043 to LK, and funded by the Center for Alzheimer’s and Related Dementias (CARD) of the National Institutes of Health (NIH) under Award Number ZO1 AG000534-02, and funded, in part, through the Intramural Research Program of the National Institutes of Neurological Disorders and Stroke (MEW). The research was also supported by an NSF Graduate Research Fellowship to OT, Tau Consortium Investigator Awards (Rainwater Charitable Foundation) to JKI, LG and MK, a Chan Zuckerberg Initiative Ben Barres Early Career Acceleration Award to MK. MK. is a Chan Zuckerberg Biohub Investigator. Development of new features for CRISPRbrain was supported, in part, by a collaboration among the Kampmann Lab, UCSF and Data Tecnica International, LLC and funding from the Center for Alzheimer’s and Related Dementias, National Institutes of Health.

## AUTHOR CONTRIBUTIONS

The iTF-Microglia differentiation strategy was developed and characterized by CH and LG with contributions from CC, LZ, JCO, LK, JI and MW. Additionally, ND, SS, OT, KL, JH and MK contributed to the optimization and characterization of iTF-Microglia. CRISPR-based functional genomics studies were designed, conducted and analyzed by ND, SS and MK with contributions from OT, KL, GA and JH. SS led the computational analysis of all screens and RNA-Seq experiments. SHH and FF developed new features for the CRISPRbrain data commons with critical input from SS, ND and MK and feedback from MAN and ABS. ND, SS, CH, OT and MK created the Figures. ND, SS and MK conceptualized and wrote the manuscript with input from the other authors. All authors reviewed and approved the final manuscript.

## METHODS

### Human iPSC culture

Human iPSCs (male WTC11 background, PMID 24509632) were cultured in StemFlex™ Basal Medium (Gibco; Cat. No. A33493-01) on BioLite Cell Culture Treated Dishes (Thermo Fisher Scientific; assorted Cat. No.) coated with Growth Factor Reduced, Phenol Red-Free, LDEV-Free Matrigel Basement Membrane Matrix (Corning; Cat. No. 356231) diluted 1:100 in Knockout DMEM (GIBCO/Thermo Fisher Scientific; Cat. No. 10829-018). StemFlex was replaced every other day or every day once 50% confluent. When 70%-80% confluent, cells were passaged by aspirating media, washing with DPBS (Gibco; Cat. No. 14190-144), incubating with StemPro Accutase Cell Dissociation Reagent (GIBCO/Thermo Fisher Scientific; Cat. No. A11105-01) at 37°C for 7 min, diluting Accutase 1:5 in StemFlex, collecting cells in conicals, centrifuging at 220g for 5 min, aspirating supernatant, resuspending cell pellet in StemFlex supplemented with 10 nM Y-27632 dihydrochloride ROCK inhibitor (Tocris; Cat. No. 125410), counting, and plating onto matrigel-coated plates at the desired number. Human iPSCs studies at the University of California, San Francisco (UCSF) were approved by the Human Gamete, Embryo and Stem Cell Research Committee.

### Human CRISPR iTF-iPS cell line generation

The two donor plasmids for inducible expression of six codon-optimized transcription factors were constructed using the plasmid pUCM (GENEWIZ). Human iPSCs (WTC11), acquired from Dr. Bruce Conklin (Gladstone Institute, San Francisco), were engineered to express PU.1, CEBPβ, and IRF5 under a doxycyline-inducible system in the CLYBL safe harbor locus and MAFB, CEBPα, and IRF8 in the AAVS1 safe harbor locus using TALEN-based editing as previously described^15^. Clones were selected using both neomycin and puromycin, thus generating the cell line we termed iTF-iPSCs. Next, iTF-iPSCs were transfected with either pC13N-dCas9-BFP-KRAB^15^, pRT029-CLYBL-CAG-DHFR-dCas9-BFP-KRAB-NLS-DHFR^15^, or pRT043-CLYBL-DDdCas9VPH-GFP^16^ to generate constitutive CRISPRi, inducible CRISPRi, or inducible CRISPRa iTF-iPS cell lines, respectively, in the CLYBL safe harbor locus using the same TALEN-editing method. After transfection, BFP-positive (CRISPRi) or GFP-positive (CRISPRa) iTF-iPSCs were repeatedly enriched via FACS sorting (BD FACS Aria Fusion). To generate monoclonal cell lines, 5,000 polyclonal CRISPR-iTF-iPSCs were plated on 10-cm dishes to enable isolation of individual clones under direct visualization with an inverted microscope (Evos FL, Thermo Fisher Scientific) in a tissue culture hood via manual scraping. Monoclonal cell lines were tested for iTF-Microglia differentiation capability and CRISPRi/a activity.

### Human iPSC-derived iTF-Microglia cell culture and differentiation

iTF-iPSCs were grown in StemFlex until reaching at least 50% confluency and were grown for at least 24h without ROCK inhibitor. They were dissociated and centrifuged as described above and pelleted cells were resuspended in Day 0 differentiation medium containing the following: Essential 8™ Basal Medium (Gibco; Cat. No. A15169-01) as a base, 10 nM ROCK inhibitor, and 2 μg/ml Doxycycline (Clontech; Cat. No. 631311). iTF-iPSCs were counted and seeded onto double coated plates (Poly-D-Lysine-precoated Bio plates (Corning, assorted Cat. No.) + Matrigel coating) with the following seeding densities: 10,000 cells/well for 96-well plate, 0.1 million/well for 12-well plate, 0.15 million/well for 6-well plate, 2 million/dish for 10-cm dish, and 8 million/dish for 15-cm dish. On day 2, media was replaced with Day 2 differentiation media containing Advanced DMEM/F12 Medium (Gibco; Cat. No. 12634-010) as a base medium containing the following: 1X Antibiotic-Antimycotic (Anti-Anti) (Gibco; Cat. No. 15240-062), 1X GlutaMAX™ (Gibco; Cat. No. 35050-061), 2 μg/ml doxycycline, 100 ng/ml Human IL34 (Peprotech; Cat. No. 200-34) and 10 ng/ml Human GM-CSF (Peprotech; Cat. No. 300-03). Two days later, on day 4, the medium was replaced with iTF-Microglia medium, containing Advanced DMEM/F12 as a base medium and the following: 1X Anti-Anti, 1X GlutaMAX, 2 μg/ml doxycycline, 100 ng/ml Human IL-34 and 10 ng/ml Human GM-CSF, 50 ng/ml Human M-CSF (Peprotech; Cat. No. 300-25) and 50 ng/ml Human TGFB1 (Peprotech; Cat. No. 100-21C). On Day 8, the media was replaced with fresh iTF-Microglia medium. iTF-Microglia can be cultured for at least 12 more days in iTF-Microglia medium with full medium changes every 3-4 days. Cells were assayed on day 8, day 9 or day 15 in most experiments. When differentiating the inducible CRISPRi/a iTF-Microglia, the media was supplemented with 50 nM trimethoprim (MP Biomedical, LLC; Cat. No. 195527) and changed every two days to maintain strong knockdown/overexpression. For dissociation, iTF-Microglia were washed once with PBS before adding TrypLE Express (Gibco; Cat. No. 12605-028) and incubating for 10 min at 37 °C. Cells were diluted 1:3 in Advanced DMEM/F12 and spun down at 220g for 5 min before resuspending in appropriate media.

### Doxycycline removal assay after Day 8 of differentiation

10,000 iTF-iPSCs were seeded into 96-well Flat Clear Bottom White Polystyrene Poly-D-Lysine Coated Microplates (Corning; Cat. No. 3843) and differentiated into iTF-Microglia as described above. At Day 8 of the differentiation, the media of iTF-Microglia was replaced with either i) full media change of iTF-Microglia medium containing 2 μg/ml doxycycline, or ii) full media change of iTF-Microglia medium containing no doxycycline, or iii) half media changes of iTF-Microglia medium containing no doxycycline. This media-replacing paradigm was repeated every three days until Day 15. Microglia survival was assessed by performing the CellTiter Glo 2.0 (Promega; Cat. No. G9242) assay according to the manufacturer’s instructions. Luminescence signal was recorded with the M5 plate reader (SpectroMax).

### Differentiation and culture of iPSC-derived microglia following the protocol by Brownjohn and colleagues

Brownjohn iPSC-Microglia (Brownjohn-iMG) were differentiated from dCas9-KRAB iPSCs (AICS-0090, Allen Cell Collection) using the published protocol^22^ with minor modifications. In brief, iPSCs (cultured in Stem Flex media with colonies at 60-80% confluency) were dissociated to single cells with Accutase, collected and plated at 10,000 cells per well in 96-well ultra-low attachment, round bottom plates (Corning; Cat. No. 7007) in 100 μl embryoid body medium (10 mM ROCK inhibitor, 50 ng/mL BMP-4 (Peprotech; Cat. No. 120-05), 20 ng/mL SCF (Peprotech; Cat. No. 300-07), and 50 ng/mL VEGF (Peprotech; Cat. No. 100-20) in E8 medium), and then subjected to centrifugation at 300g for 3 min. Embryoid bodies were cultured for 4 days, with a half medium change after 2 days. On day 4, embryoid bodies were carefully collected and transferred into a 15 ml conical tube, and left to settle at the bottom. The embryoid media was aspirated and 15 to 20 embryoid bodies were plated per well in 6-well plates and cultured in 3 mL hematopoetic medium (2 mM GlutaMax, 1x Anti-Anti, 55 mM 2-mercaptoethanol (BioRad; Cat. No. 1610710), 100 ng/mL M-CSF, and 25 ng/mL Human IL-3 (Peprotech; Cat. No. 200-03) in X-Vivo 15 (Lonza; Cat. No. BE02-060F). Two thirds of the media was exchanged every 3-4 days. Microglia progenitors were harvested from suspension after 14-21 days and plated onto PDL-coated plates in microglia maturation media (2 mM GlutaMax, 1x Anti-Anti, 100 ng/mL IL-34, and 10 ng/mL GM-CSF in Advanced RPMI-1640 (Gibco; Cat. No. 12633012)). Microglia progenitors were further differentiated for 8 days with full medium change every 2–3 days before using them for experiments.

### iTF-Microglia coculture with iNeurons

iPSC-derived neurons (iNeurons) were differentiated from WTC11 iPSCs engineered to express NGN2 under a doxycycline-inducible system in the AAVS1 safe harbor locus as previously described^15, 84^ with minor modifications as follows: iPSCs were maintained and dissociated as described above and replated on Matrigel coated dishes in N2 Pre-Differentiation Medium. After three days, hereafter Day 0, the pre-differentiated neurons were dissociated to single cells with Accutase, collected, and plated at 10,000 cells per well in PDL-coated 96-well plates in BrainPhys Neuronal Medium (BrainPhys (STEMCELL Technologies; Cat. No. 05790) as the base, 0.5x N2 supplement (Thermo Fisher; Cat. No. 17502-048), 0.5x B27 Supplement (GIBCO/Thermo Fisher Scientific; Cat. No. 17504-044), 10ng/mL NT-3 (PeproTech; Cat. No. 450-03), 10ng/mL BDNF, 1mg/mL Mouse laminin (Thermo Fisher; Cat. No. 23017-015), and 2mg/mL doxycycline. On Day 3, a full media change was performed. On day 7, half the media was removed, and an equal volume of BrainPhys Neuronal Medium was added. On day 14, half the media was removed and an equal volume of BrainPhys Neuronal Medium containing Day 8 iTF-Microglia expressing Lck-mNeonGreen and supplemented with 2X the cytokines of the iTF-Microglia medium was added. 3,000 iTF-Microglia were added to each well and immunostaining experiments were performed after one day.

### Lentiviral transduction of iPSCs with sgRNA constructs

Individual or pooled sgRNAs were introduced into CRISPRi- or CRISPRa-iPSCs via lentiviral delivery using TransIT Lenti Reagent (Mirus Bio LLC; Cat. No. MIR 6600) according to manufacturer’s protocol. Cells were selected with 2 µg/ml Puromycin (Gibco; Cat. No. A11138-03) for 2 - 4 days and recovered 2 - 4 days until MOI >0.9 as determined by flow cytometry of sgRNA-BFP fluorescence. sgRNA protospacer sequences are provided in Supplementary Table 10.

### qPCR

To quantify *TFRC, INPP5D or PICALM* knockdown or *CXCR4, CD209 or PFN1* overexpression, lysed cell pellets from human iPSCs or iTF-Microglia were thawed on ice, and total RNA was extracted using the Quick-RNA Miniprep Kit (Zymo; Cat. No. R1054). cDNA was synthesized with the SensiFAST cDNA Synthesis Kit (Bioline; Cat. No. 65054). Samples were prepared for qPCR in technical triplicates in 5 µL reaction volumes using SensiFAST SYBR Lo-ROX 2X master mix (Bioline; Cat. No. BIO-94005), custom qPCR primers from Integrated DNA Technologies used at a final concentration of 0.2 µM, and cDNA diluted at 1:3. Quantitative real-time PCR was performed on an Applied Biosystems QuantStudio 6 Pro Real-Time PCR System with the following Fast 2-Step protocol: (1) 95° C for 20 s; (2) 95° C for 5 s (denaturation); (3) 60°C for 20 s (annealing/extension); (4) repeat steps 2 and 3 for a total of 40 cycles; (5) 95°C for 1 s; (6) ramp 1.92°C/s from 60°C to 95°C to establish melting curve. Expression fold changes were calculated using the ΔΔCt method normalizing to housekeeping gene *GAPDH*. Primer sequences are provided in Supplementary Table 10.

### Cell surface protein staining for flow cytometry

Dissociated and resuspended iTF-Microglia were blocked for 15 min with 1:20 Human FC Block (BD Biosciences; Cat. No. 564220) and then stained with 1:66 PE/Cy7 anti-human CD184 (CXCR4) (BioLegend; Cat. No. 306514) for CRISPRa validation or 1:66 PE-Cy7 anti-human CD71 (TFRC) (BioLegend; Cat. No. 334112) for CRISPRi validation for 30 min in the dark. For the CD38 screen and validation experiments, iTF-Microglia were stained with 1:200 anti-hCD38 PE (R&D Systems; Cat. No. FAB2404P).

Cells were washed twice with DPBS before analyzing them by flow cytometry using the BD LSRFortessa X14 (BD Biosciences). Flow cytometry data was analyzed using FlowJo (FlowJo, version 10.7.1), raw median fluorescence intensity values of CD184, CD71 and CD38 stained cells were normalized to non-stained control samples and data was plotted as fold change using Prism 8 (GraphPad, version 8.4.2).

### Intracellular protein staining for flow cytometry

iTF-Microglia were treated for 6 h with 1:2000 GolgiPlug™ (BD; Cat. No. 555029) or DMSO as control before dissociating. Cells were fixed and permeabilized with the eBioscience Intracellular Fixation and Permeabilization Buffer Set (Invitrogen; Cat. No. 88-8824-00) according to the manufacturer’s instructions. Cells were stained with 1:75 Anti-Hu Osteopontin (SPP1) eFluor 660 (eBioscience; Cat. No. 50-9096-42) or 1:75 Human CCL13 488 (R&D Systems; Cat. No. IC327G) or their isotype controls Mouse IgG1 Control Alexa Fluor 488 conjugated (R&D Systems; Cat. No. IC002G) and Mouse IGG1 kappa Isotype (eBioscience; Cat. No. 50-4714-82) over night at 4 °C. Cells were washed twice with DPBS before analyzing them by flow cytometry using the BD FACS CelestaTM (BD Biosciences) or the BD FACSAria Fusion. Flow cytometry data was analyzed using FlowJo, raw median fluorescence intensity values of Osteopontin (SPP1) and CCL13 stained cells were normalized to isotype-control samples and data was plotted as fold change using Prism 8. The gating strategy used to determine the percentage of SPP1-positive cells is shown in Supplementary Fig. 1c).

### Immunohistochemistry

iTF-Microglia monocultures and co-cultures were differentiated in PDL-coated 96-well plates. They were fixed with 4% Paraformaldehyde (Electron Microscopy Sciences; Cat. No. 15710) for 10 min at room temperature. After washing with DPBS 3 times, cells were permeabilized and blocked with 5% normal goat serum (Vector Laboratories; Cat. No. S-1000-20) with 0.01% Triton X-100 (TEKnova; Cat. No. T1105) in PBS for 1 hr at room temperature. Cells were then incubated with primary antibodies diluted in blocking buffer at 4 °C overnight. After that, cells were washed with DPBS 3 times and incubated with secondary antibodies diluted in blocking buffer for 1 hr at room temperature. Cells were then washed with DPBS 3 times and stained with 10 μg/ml Hoechst 33342 (Thermo Fisher Scientific; Cat. No. H3570) for 10 min. Cells were imaged using a confocal microscope (Leica SP8) or an IN Cell Analyzer 6000 (GE; Cat. No. 28-9938-51). Primary antibodies used for immunofluorescence in this study were as follows: anti-mouse 1:150 GPR34 (R&D Systems; Cat. No. MAB4617), anti-rabbit 1:1000 IBA1 (Wako; Cat. No. 019-19741), anti-rabbit 1:200 TFRC (abcam; Cat. No. ab84036), anti-rabbit 1:1000 synaptophysin (Synaptic Systems; Cat. No. 101 004). Secondary antibodies used in this study were as follows: goat anti-rabbit IgG Alexa Fluor 555 (1:500 dilution; abcam; Cat. No. ab150078), goat anti-mouse IgG Alexa Fluor 488 (1:500 dilution; abcam; Cat. No. ab150113) and goat anti-chicken IgG Alexa Fluor 647. F-actin was stained using ActinGreenTM 488 (Invitrogen; Cat. No. R37110) according to the manufacturer’s protocol.

### Synaptosome isolation and pHrodoRed labelling

Synaptosomes were isolated from fresh Innovative Grade US Origin Rat Sprague Dawley Brain (Innovative Research, Inc.; Cat. No. IGRTSDBR) with the Syn-PER™ Synaptic Protein Extraction Reagent (Thermo Scientific™; Cat. No. 87793) according to the manufacture’s protocol with minor changes. Briefly, 10 mL of Syn-PER Reagent supplemented with 1x protease inhibitor cOmplete Mini, EDTA free (Roche; Cat. No. 11836170001) and 1x phosphatase inhibitor PhosSTOP (Roche; Cat. No. 4906845001) were added per gram of brain tissue. Dounce homogenization was performed on ice and homogenate was transferred to a conical tube and centrifuged at 1200 × g for 10 minutes at 4°C. The pellet was discarded, the supernatant was transferred to a new tube, and the centrifugation step was repeated. The supernatant was then centrifuged at 15,000 × g for 20 minutes at 4°C. The supernatant was removed and the wet pellet was weighed. The synaptosome fractions were resuspended at a concentration of 50 mg/ml. 3 μM pHrodo™ Red, succinimidyl ester (pHrodo™ Red, SE) (ThermoFisher Scientific; Cat. No. P36600) was added to the synaptosome fraction and incubated for 45 min at room temperature in the dark. After diluting the solution 1:10 in DPBS, the synaptosomes were spun down at 2500 × g for 5 min. The supernatant was removed and then the synaptosomes were washed two times with DPBS. The pHrodo-labeled synaptosomes were resuspended in microglia iTF-Microglia medium at a stock concentration of 50 mg/ml and directly used for phagocytosis assays or frozen in synaptosome freezing media (5% DMSO in Advanced DMEM/F12) for later use.

### Phagocytosis assays

Day 8 iTF-Microglia were used for all phagocytosis assays. iTF-Microglia medium was prepared with pHRodoRed-labeled synaptosomes at a concentration of 1 mg/ml or 0.5 μl/ml media of Fluoresbrite Carboxylate YG 1.0 Micron Microspheres (15702-10; Cat. No. 15702-10). After replacing the media with the substrate media, iTF-Microglia were incubated for 1.5 h in the incubator if not otherwise stated. Cells were washed twice with DPBS, dissociated, resuspended in ice-cold DPBS, and analyzed via flow cytometry. Where indicated, actin polymerization was inhibited by pretreating cells with 5 μM Cytochalasin D (Invitrogen; Cat. No. PHZ1063) for 30 min before the addition of phagocytic substrate media. For analyzing phagocytic capabilities within microglia clusters, pHRodoRed-labeled synaptosomes at a concentration of 1 mg/ml were added to iTF-Microglia for 1.5h. Microglia were washed 3x with PBS before incubating them in iTF-Microglia media supplemented with 1:2000 GolgiPlug™ (BD; Cat. No. 555029) for 4h. Cells were dissociated, fixed, and stained for CCL13 and SPP1 as described above. Flow cytometry data was analyzed using FlowJo, raw median fluorescence intensity values of phagocytosing cells were normalized to no-substrate control samples and data plotted as fold change using Prism 8.

### Human cytokine array

Day 8 iTF-Microglia were treated with 100 ng/ml LPS (Millipore Sigma; Cat. No. LPS25) or DPBS control. After 24 hours, the supernatant was collected and processed using the Proteome Profiler Human Cytokine Array Kit (R&D Systems; Cat. No. ARY005B), according to manufacturer’s instructions. For analysis of the signals, Fiji (version 2.0.0) was used to measure the integrated pixel density for each pair of duplicate dots representing a cytokine. Background signal was measured from negative control dots and then subtracted from each dot. The relative change in cytokine levels as a result of LPS-treatment was obtained by comparing corresponding cytokine signals across multiple arrays performed in tandem.

### Live-cell imaging

iTF-iPSCs transduced with individual sgRNAs as described above were passaged and differentiated into iTF-Microglia in the 96-well format described above. Starting on day 2 of differentiation, and continuing every two days until day 15, iTF-Microglia were stained with 10 μg/ml Hoechst 33342 for 10 min at 37 °C, washed with PBS, and imaged with the IN Cell Analyzer 6000. Using the same 96-well format as described above, Day 8 iTF-Microglia were stained for F-actin using 25 nM SiR-actin (Cytoskeleton, Inc; Cat. No. CY-SC001) probe, diluted in iTF-Microglia medium, with a 4 hour incubation at 37 °C. A full media change with iTF-Microglia medium was completed before imaging using the IN Cell Analyzer 6000.

### CellTiter Glo assay after pharmaceutical inhibition of CSF1R or MAPK14

10,000 iTF-iPSCs were seeded into 96-well Flat Clear Bottom White Polystyrene Poly-D-Lysine Coated Microplates (Corning; Cat. No. 3843) and differentiated into iTF-Microglia. For CSF1R inhibition, cells were treated with the CSF1R inhibitor PLX3397 at Day 8 (ApexBio; Cat. No. B5854) at indicated concentrations or DMSO control for 24h before performing the CellTiter Glo 2.0 (Promega; Cat. No. G9242) assay according to the manufacturer’s instructions. For MAPK14 inhibition, cells were treated with the MAPK14 inhibitor Skepinone-L at Day 8 (Selleckchem; Cat. No. 1221485831) at indicated concentrations or DMSO control for 24h before performing the CellTiter Glo 2.0 assay. Luminescence signal was recorded with the M5 plate reader (SpectroMax).

## CRISPR SCREENS

### Large-scale survival-based and FACS-based screens

Inducible CRISPRi and CRISPRa iTF-iPSCs were infected with pooled CRISPRi or CRISPRa sgRNA libraries^33^ targeting the druggable genome and selected for lentiviral integration with puromycin, as described above. Day 0 iTF-iPSCs, with a cell count corresponding to an average 1000x coverage per library element, were differentiated into iTF-Microglia as described above with constant TMP supplementation for dCas9 stabilization.

For the survival screens, Day 0 iPSCs and Day 15 iTF-Microglia were lifted with Accutase or TryplE Express, respectively. Lifted cells were harvested and subjected to sample preparation for next-generation sequencing as described below.

For the CD38-activation screen, Day 8 iTF-Microglia were dissociated with TrypleE and then blocked and stained with anti-PE-CD38 as described in the cell surface staining section. Cells were sorted into high and low signal population corresponding to the top 30% and the bottom 30% of the CD38-PE signal distribution (gating strategy shown in Supplementary Fig. 1a).

For the phagocytosis FACS screen, Day 15 iTF-Microglia were incubated with PhRodo-Red synaptosomes as described in phagocytosis assay section. Cells were then dissociated with TryplE and sorted into high and low signal population corresponding to the top 30% and the bottom 30% of the PhRodoRed signal distribution (gating strategy shown in Supplementary Fig. 1b). Based on simulations, we previously found that this sorting strategy is optimal for hit detection in FACS-based screens^85^.

Cells were subjected to sample preparation for next-generation sequencing as previously described^15^. Briefly, for each screen sample, genomic DNA was isolated using a Macherey-Nagel Blood L kit (Macherey-Nagel; Cat. No. 740954.20). sgRNA-encoding regions were amplified and sequenced on an Illumina HiSeq-4000.

### Quant-Seq

Cell culture medium was aspirated, cells were washed once with DPBS, and RNA lysis buffer was added directly to wells containing either Day 0 iTF-iPSCs, Day 15 iTF-Microglia +/- 50 ng/ml 24h LPS treatment, Brownjohn-iMG +/- 100 ng/ml 24h LPS treatment, or Day 15 iTF-Microglia. For assessing transcriptomic effects after *PFN1* overexpression, two different *PFN1* sgRNAs and NTC sgRNAs were transduced into inducible CRISPRa iPSCs and cells were differentiated to Day 8 iTF-Microglia. Biological triplicates for each condition (approximately 0.15 Mio cells each) were pelleted, snap frozen, and stored at -80°C. RNA was extracted using the Quick-RNA Miniprep Kit (Zymo; Cat. No. R1055). Libraries were prepared from total RNA (250-473 ng per sample) using the QuantSeq 3‘ mRNA-Seq Library Prep Kit for Illumina (FWD) (Lexogen; Cat. No. 015UG009V0252) following the manufacturer’s instructions. Library amplification was performed with 14 total PCR cycles. mRNA-seq library concentrations (mean of 1.13 ± 0.66 ng/uL) were measured with the Qubit dsDNA HS Assay Kit (Invitrogen; Cat. No. Q32851) on a Qubit 2.0 Fluorometer. Library fragment-length distributions (mean of 287 ± 28 bp) were quantified with High Sensitivity D5000 Reagents (Agilent Technologies; Cat. No. 5067-5593) on the 4200 TapeStation System. The libraries were sequenced on an Illumina NextSeq 2000 instrument with single-end reads.

### CROP-Seq

A pooled sgRNA library consisting of 2 sgRNAs per targeted gene and 4 non-targeting control sgRNAs was designed to target 39 genes which were selected hit genes from iTF-Microglia survival and FACS-based screens (Supplementary Table 5; only 1 sgRNA for gene *DBF4* due to technical error). Briefly, top and bottom strands of sgRNA oligos were synthesized (Integrated DNA Technologies) and annealed in an arrayed format, pooled in equal amounts, and ligated into our optimized CROP-seq vector, as previously described^15^.

Inducible CRISPRi-iTF-iPSCs were infected with the pooled sgRNA library at <0.15 MOI and then selected for lentiviral integration. Next, iTF-iPSCs were differentiated into iTF-microglia and cultured with the addition of TMP. Day 8 iTF-Microglia were washed 3X with DPBS, dissociated with TrypLE, and resuspended in nuclease-free water before loading onto four wells of the 10x Chromium Controller (10x Genomics, v3.1) according to the manufacturer’s protocol, with 35,000 cells recovered per sample as the target. Sample preparation was performed using the Chromium Next GEM Single Cell 3′ Reagent Kits version 3.1 (10x Genomics, cat. no. PN-1000121) according to the manufacturer’s protocol, reserving 10-30 ng full-length cDNA to facilitate sgRNA assignment by amplifying sgRNA-containing transcripts using hemi-nested PCR reactions adapted from a previously published approach^15, 86^. cDNA fragment analysis was performed using the 4200 TapeStation System and sgRNA enrichment libraries were separately indexed and sequenced as spike-ins alongside the whole-transcriptome scRNA-seq libraries using a NovaSeq 6000 using the following configuration: Read 1: 28; i7 index: 8; i5 index: 0; Read 2: 91.

## COMPUTATIONAL AND STATISTICAL ANALYSIS

### Primary CRISPR screen analysis

Primary screens were analyzed using our previously published MAGeCK-iNC bioinformatics pipeline^15^, available at https://kampmannlab.ucsf.edu/mageck-inc. Briefly, raw sequencing reads from next-generation sequencing were cropped and aligned to the reference using Bowtie version 0.12.9^87^ to determine sgRNA counts in each sample. The quality of each screen was assessed by plotting the log10(counts) per sgRNA on a rank order plot using ggplot2 version 3.3.3^88^. Raw phenotype scores and significance P values were calculated for target genes, as well as for ‘negative-control-quasi-genes’ that were generated by random sampling with replacement of five non-targeting control sgRNAs from all non-targeting control sgRNAs. The final phenotype score for each gene was calculated by subtracting the raw phenotype score by the median raw phenotype score of ‘negative-control-quasi-genes’ and then dividing by the standard deviation of raw phenotype scores of ‘negative-control-quasi-genes’.

A ‘Gene Score’ was defined as the product of phenotype score and –log10(P value). Hit genes were determined based on the Gene Score cutoff corresponding to an empirical FDR of 10%. Volcano plots of gene scores were generated using ggplot2 version 3.3.3^88^.

### RNA-seq analysis

Alignment and mapping were performed using Salmon version 1.4.0^89^, (the --noLengthCorrection flag was used for QuantSeq samples) and either the human reference genome GRCh38 (Gencode, release 37), or a custom GRCh38 reference genome containing the references for each 3TF transgene integrated in iTF-iPSCs, to obtain transcript abundance counts. Tximport version 1.18.0^90^ was used to obtain gene-level count estimates. Genes with zero counts across all samples were removed from the analysis. To visualize differences in gene expression across samples, a list of gene symbols corresponding to microglia markers, microglia activation markers, and iPSC markers were compiled from a previous publication^91^; the normalized counts of each of these genes were then standardized across samples (*i.e.* subtracting by the mean and dividing by the standard deviation) and visualized using Complex Heatmap version 2.6.2^92^. To assess how iTF-Microglia compare to other iPSC-Microglia and primary microglia from a range of previous studies^17, 22, 24, 25, 93^, raw fastqs obtained from the NCBI GEO database) were subject to the same analysis pipeline stated above. Then, principal component analysis was performed using microglia marker genes as input with DESeq2 version 1.30.1^94^.

For differential gene expression analysis of LPS-treated vs. PBS-treated iTF-Microglia samples and the *PFN1* overexpression vs. NTC iTF-Microglia DESeq2 version 1.30.1^94^ was used to calculate the log-fold-change and p-values and perform shrinkage of log-fold-change for downstream visualization using ggplot2 version 3.3.3^88^. To compare both LPS-treatment up-regulated and down-regulated differentially expressed genes (DEGs) in iTF-Microglia and Brownjohn-iMG, DEGs that were significant (padj < 0.05) in at least one cell type were visualized using VennDiagram (version 1.6.20).

### CROP-seq analysis

Alignment and gene expression quantification was performed on scRNAseq libraries and sgRNA-enriched libraries using Cell Ranger (version 5.0.1, 10X Genomics) with default parameters and reference genome GRCh38-3.0.0. Cellranger aggr was used to aggregate counts belonging to the same sample across different GEM wells. The resulting gene vs. cell barcode matrix contained 58,302 cells which had on average 41,827 reads per cell, and a median of 3,346 genes per cell. sgRNA unique molecular identifier (UMI) counts for each cell barcode were obtained using a previously described mapping workflow^86^. To facilitate sgRNA identity assignment, a combination of demuxEM ^95^ and a z-score cut-off method we previously described^15^ were used such that only cells with a single sgRNA as determined by both methods were carried forward in the analysis.

The raw gene vs. barcode matrix outputted by Cell Ranger was converted into a SingleCellExperiment (SCE) object using the read10xCounts function from the DropletUtils package version 1.10.3^96^ in R (version 4.0.3). sgRNA assignments were appended to the SCE metadata and filtered to only include cells with a single sgRNA, resulting in 28,905 cells. The SCE was converted into a Seurat object using Seurat::as.Seurat version 4.0.1^97^. The data was normalized and highly variable genes were identified using Seurat::SCTransform^98^. For initial data exploration, principal-component analysis was performed using Seurat::RunPCA to determine the number of principal components to retain. UMAP dimensional reduction using Seurat::RunUMAP and clustering using Seurat::FindNeighbors and Seurat::FindClusters were performed on the retained principal components with resolution = 0.7.

Initial data exploration revealed clusters that were not of interest due to a high proportion of mitochondrial-encoding genes or disrupted microglia differentiation (Extended Data Fig.5A). These clusters were removed from the downstream analysis and the remaining “microglia cluster” population was normalized, clustered, and visualized using UMAP, as described above with resolution = 0.25.

To determine the differentially expressed genes between UMAP clusters, Seurat::FindAllMarkers was used. Single-cell heatmaps, ridge plots, rank plots, and UMAPs were made using Seurat::DoHeatmap, Seurat::RidgePlot, Seurat::VizDimLoadings, and Seurat::DimPlot or Seurat::FeaturePlot, respectively.

The relative proportion of cells with a given sgRNA in a given cluster compared to cells with non-targeting control (NTC) sgRNAs in the given cluster was calculated and visualized using Complex Heatmap version 2.6.2^92^ (Supplemental Table 7).

For each CRISPRi target gene, the population of cells with the strongest knockdown (cells with expression of target gene less than the median expression of the target gene) was carried forward to perform differential gene expression analysis using Seurat::FindMarkers with parameters test.use = ‘t’ (student’s t-test), assay = “SCT”, slot = “scale.data”, to compare the Pearson residuals of cells^98^ with knockdown sgRNAs versus non-targeting control cells. Genes with an adjusted p-value < 0.1 were deemed significant. The top 20 DEGs for each target gene which had >50 cells comprised the set of genes used to visualize convergent pathways using Complex Heatmap version 2.6.2^92^.

The iTF-Microglia CROP-seq dataset was integrated with the previously published human scRNAseq dataset (Olah-hMG)^7^ using Seurat^99^. Briefly, the Olah-hMG gene vs. cell barcode matrix and metadata were used to create a Seurat object and cells from surgery samples or with non-microglia identities as previously determined^7^ were removed. Normalization and identification of highly variable genes was performed using Seurat::SCTransform with the same parameters as the iTF-Microglia. Next, integration features (3000 features) and integration anchors were identified for each Seurat object using Seurat::SelectIntegrationFeatures and Seurat::FindIntegrationAnchors and subsequent integration with identified anchors was performed using Seurat::IntegrateData. The integrated Seurat object was normalized, clustered, and visualized using UMAP, as described above with resolution = 0.25. Gene expression was visualized with UMAP using Seurat::FeaturePlot and the percentage of cells in the integrated SPP1-high cluster or SPP1-low clusters of either AD brain or control brain origin was calculated and significance was calculated on the cell counts using Fisher’s exact test.

### Image analysis with CellProfiler

Pipelines and example images are compiled in supplemental material and all analysis was performed using CellProfiler version 4.1.3. *Cell morphology metrics*: Nuclei were segmented as primary objects from Hoechst images. Cell segmentations were generated by propagating outward from nuclei objects until edges were identified in the phalloidin images. Area and shape metrics were calculated for each cell object. *Integrated F-actin intensity per cell*: for a given field of view, nuclei were segmented based on Hoechst images and total integrated SiR-actin intensity was summed. The resulting sum was divided and by the number of nuclei. *Longitudinal cell counts:* for a given field of view, nuclei were segmented based on Hoechst images acquired daily. *IBA1 intensity per cell:* This metric was determined similarly to the integrated F-actin intensity per nuclei; for a given field of view, the total integrated intensity of the IBA1 stain was divided by the number of segmented nuclei based on Hoechst.

### Data Availability Statement

All screen datasets and RNA-transcriptomic datasets are publicly available in the CRISPRbrain data commons (http://crisprbrain.org/) (associated with Figures 1, 2, 4, 5, 6, 7 and Extended Data Figures 1, 4, 5, 6, 7). RNA sequencing datasets reported in this paper are in the process of being deposited on NCBI GEO. There are no restrictions on data availability.

### Code Availability Statement

The MAGeCK-iNC bioinformatics pipeline for analysis of pooled screens available at https://kampmannlab.ucsf.edu/mageck-inc. The CellProfiler pipelines will be made available on request to the corresponding authors (MK), and will also be submitted to the CellProfiler depository of published pipelines (https://cellprofiler.org/examples/published_pipelines.html) upon publication.

---

**Extended Data Figure 1:**
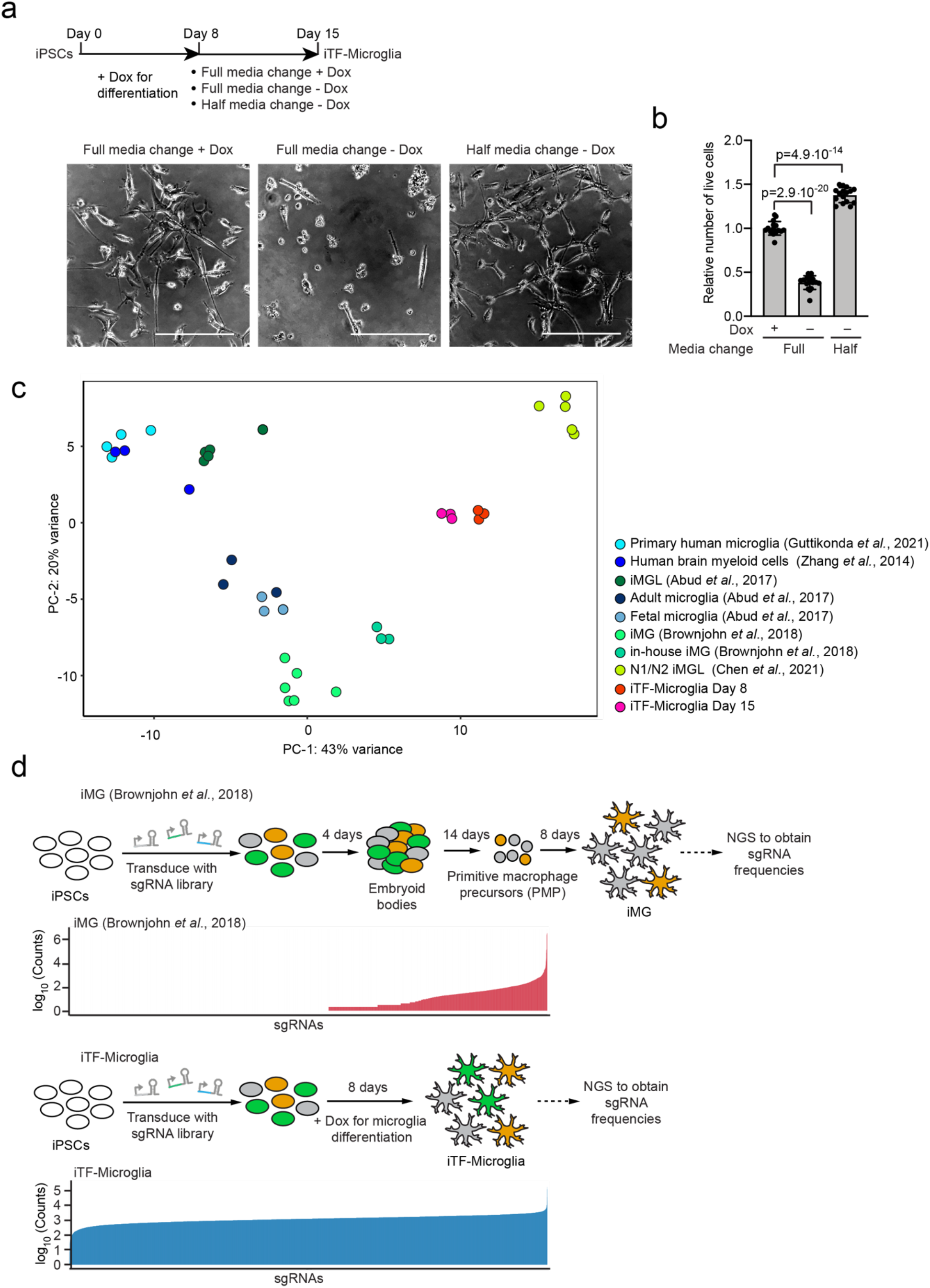
Impact of Doxycycline removal on iTF-Microglia survival and sgRNA recovery in iPSC-derived microglia generated with different protocols. **a** Comparison of iTF-Microglia viability after Day 8 with different protocols. *Top*: timeline with different doxycycline supplementation paradigms, *bottom*: representative phase-contrast images at Day 15 with the indicated doxyccycline supplementation. Scale bar: 50 μm**. b**, Survival of iTF-Microglia at Day 15 after different doxycycline treatments indicated in a. Viable cells were quantified using the CellTiter-Glo assay. Values represent mean +/- sd of n = 12 biological replicates; p values from two-tailed Student’s t-test. **c**, Principal component analysis (PCA) on the expression of microglia marker genes of iTF-Microglia, human adult ex-vivo microglia^93^, fetal and adult microglia^17^, human myeloid cells^25^, other iPSC-microglia^17, 22, 24^. No iPSC samples were included. Each dot reflects an independent biological sample. Colors represent the different cell types. **d**, sgRNA recovery after transduction with a pooled sgRNA library in iPSCs and differentiation with two different iPSC-Microglia protocols. Strategy for the infection of iPSCs with an sgRNA library with 13,025 elements and timepoint of sgRNA recovery in iPSC-Microglia with the actual recovered counts of sgRNAs after next-generation-sequencing (NGS) from the protocol from Brownjohn *et al*.^22^ (*Top*) and iTF-Microglia (*Bottom*).

**Extended Data Figure 2:**
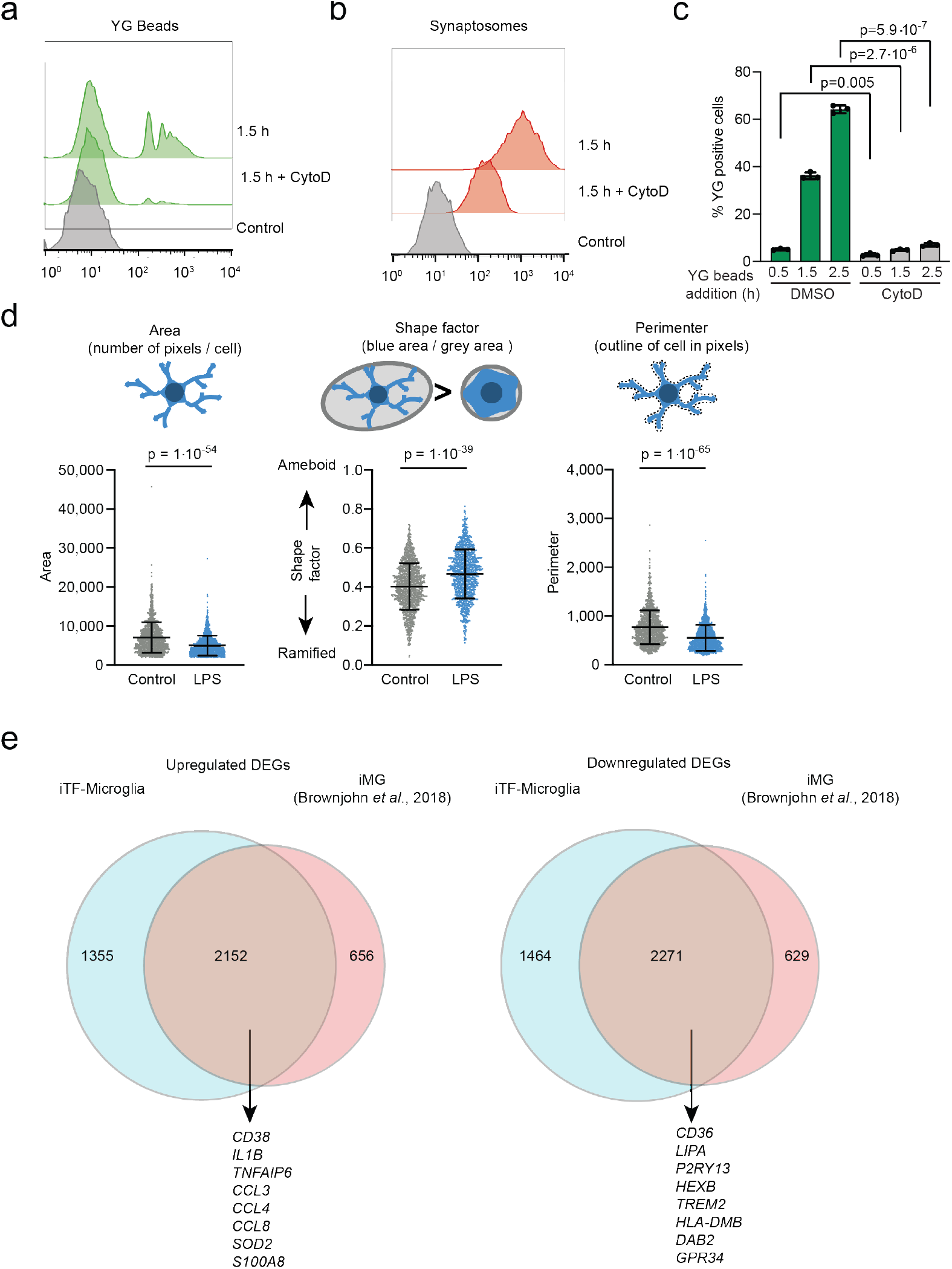
Phagocytosis capacity of iTF-Microglia and morphological changes after LPS treatment. **a-b,** Phagocytosis of yellow-green (YG) beads (a) or pHRodo-Red labelled synaptosomes (b) measured by flow cytometry. Histograms of YG-beads-FITC (a) and Synaptosome-PE (b) after 1.5h of substrate exposure +/- 5μM Cytochalasin D (CytoD) treatment. Controls are iTG-Microglia without substrate exposure. **c,** Phagocytosis of yellow-green (YG) beads at different timepoints. Flow cytometric quantification of the percentage of YG bead-positive cells at after 0.5 h, 1.5 h and 2.5 h of incubation with beads. Addition of 5 μM CytoD decreases the percentage of YG bead-positive cells. N = three individual biological replicates; p values from two-tailed Student’s t-test. **d**, Morphological changes of iTF-Microglia after LPS treatment. Swarm plots showing the automated quantification of microglia F-actin staining in area, shape factor and perimeter with explanation of the three parameters. N = 16 wells from 3 individual differentiations; p values from two-tailed Mann-Whitney test. **e,** Comparison of differentially expressed genes in response to LPS treatment in iTF-Microglia versus iPSC-derived microglia (iMG) differentiated following a previously published protocol by Brownjohn *et al*.^22^.

**Extended Data Figure 3:**
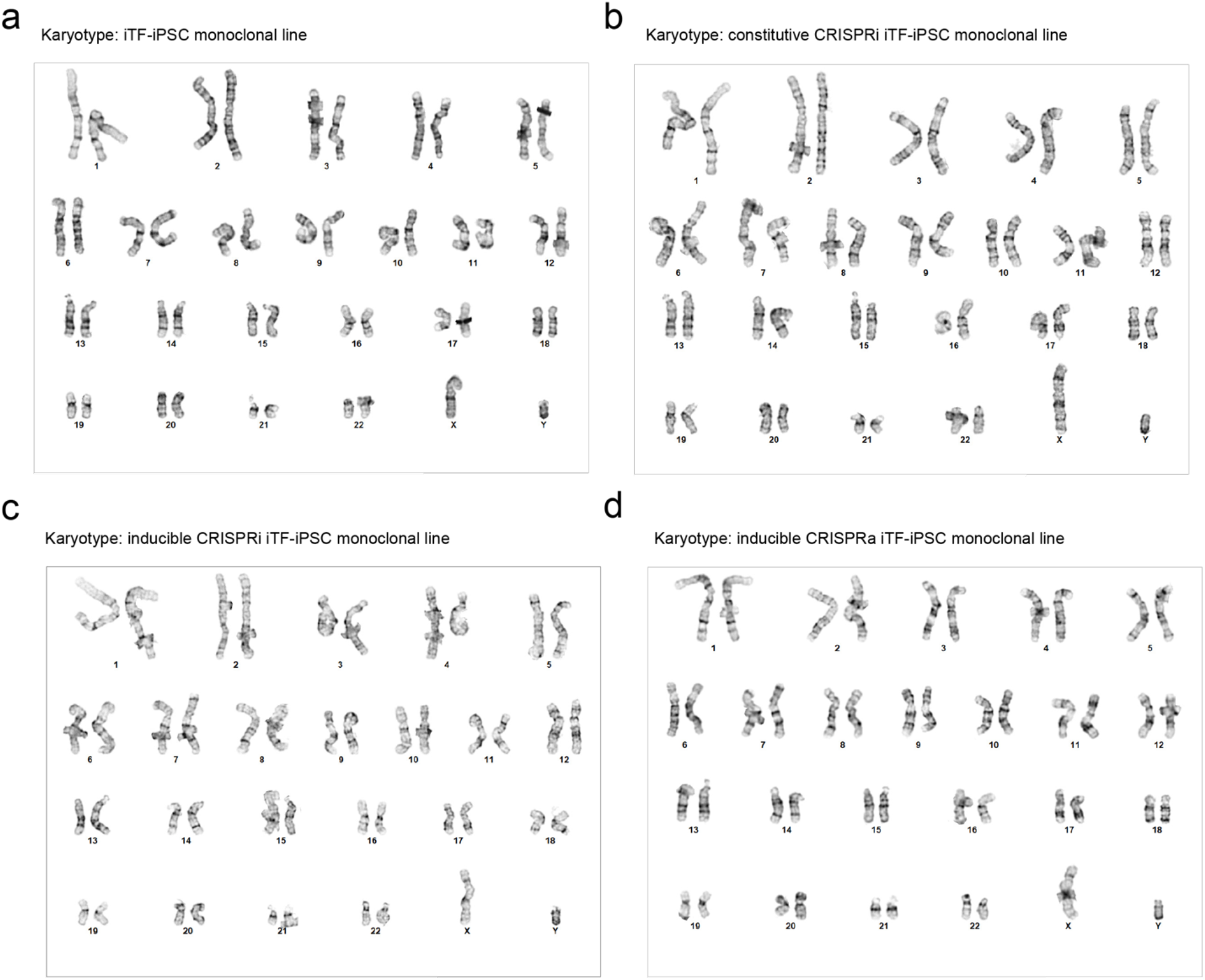
Karyotyping of the monoclonal iTF-iPSC lines. A normal karyotype was confirmed for monoclonal **a,** iTF-iPSCs**, b,** constitutive CRISPRi iTF-iPSC, **c,** inducible CRISPRi iTF-iPSC, **d,** inducible CRISPRa iTF-iPSC lines.

**Extended Data Figure 4:**
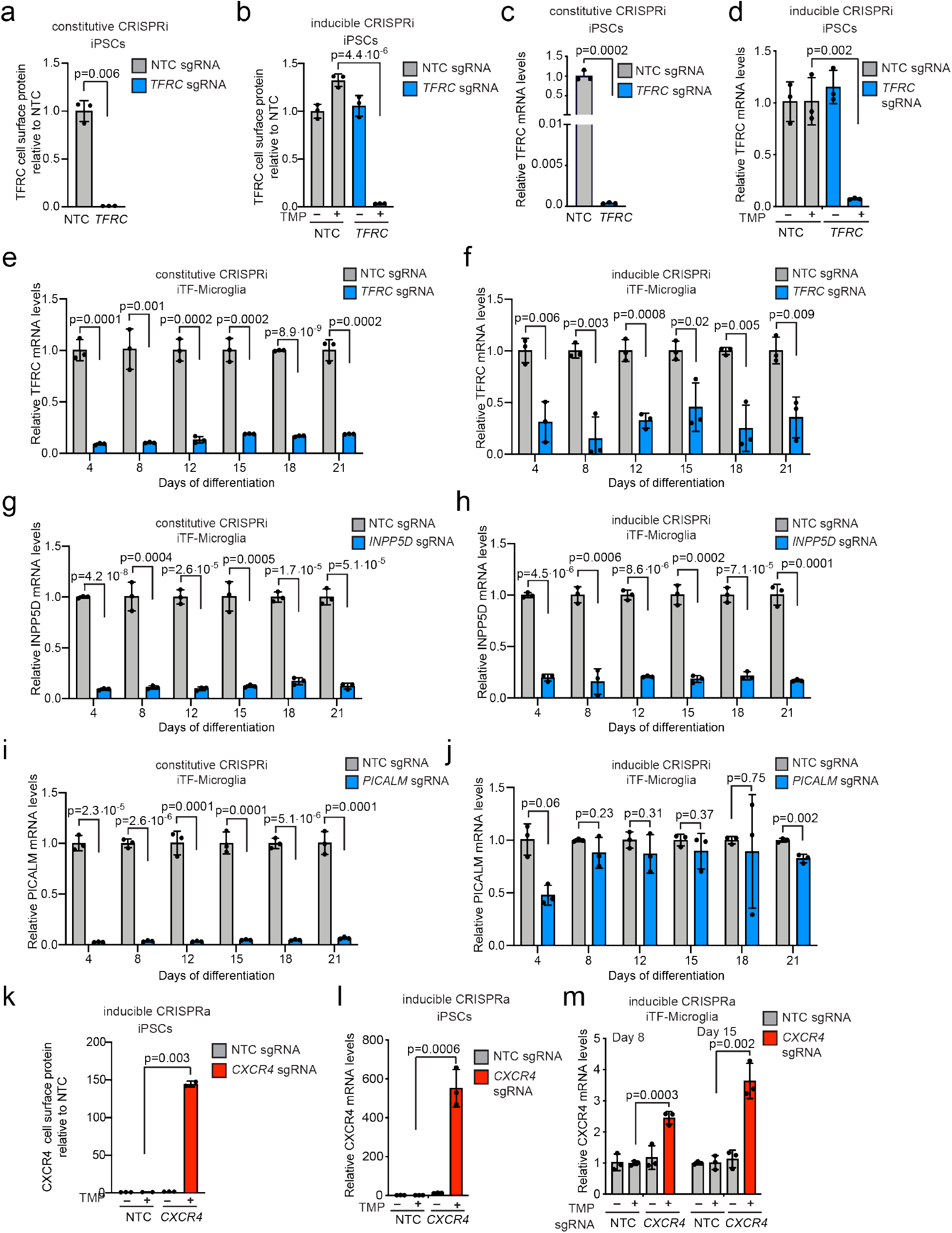
Functional validation of CRISPRi/a activity in iPSCs and iTF-Microglia. **a-b,** Functional validation of constitutive (a) or inducible (b) CRISPRi activity via flow cytometry of TFRC surface protein level stained iPSCs expressing a TFRC-targeting sgRNA or a non-targeting control (NTC) sgRNA (mean +/- sd, n = 3 biological replicates; p values from two-tailed Student’s t-test). TMP was added to induce CRISPRi activity where indicated. **c-d, Knockdown of TFRC in iPSCs with (a) the constitutive and (b) the inducible CRISPRi system.** qPCR quantification of the relative fold change of TFRC mRNA levels in CRISPRi-iPSCs expressing a TFRC sgRNA as compared to a non-targeting control sgRNA in the presence or absence of trimethoprim (TMP). (mean +/- sd, n = 3 biological replicates; p values from two-tailed Student’s t-test). TFRC levels were normalized to the housekeeping gene GAPDH. **e-j Knockdown of three different genes in iTF-Microglia with (e,g,i) constitutive CRISPRi and (f,h,j) inducible CRISPRi.** qPCR quantification of the relative fold change of *TFRC* mRNA levels (e,f), *INPP5D* mRNA levels (g,h) or *PICALM* mRNA levels (I,j) in CRISPRi-iTF-Microglia expressing a *TFRC* sgRNA (e,f), *INPP5D* sgRNA (g,h) or *PICALM* sgRNA (I,j) compared to a non-targeting control sgRNA at different days of differentiation in the presence of TMP (mean +/- sd, n = 3 biological replicates). **k,** Functional validation of inducible CRISPRa activity via flow cytometry of CXCR4 surface protein level stained iPSCs expressing a CXCR4-targeting sgRNA or a non-targeting control (NTC) sgRNA (mean +/- sd, n = 3 biological replicates; p values from two-tailed Student’s t-test). TMP was added to induce CRISPRi activity where indicated. **l-m,** qPCR quantification of the relative fold change of CXCR4 mRNA levels in inducible CRISPRa-iPSCs expressing a CXCR4 sgRNA as compared to a non-targeting control sgRNA in the presence or absence of trimethoprim (TMP), which stabilizes the DHFR degron. (mean +/- sd, n = 3 biological replicates; p values from two-tailed Student’s t-test). CXCR4 levels were normalized to the housekeeping gene GAPDH.

**Extended Data Figure 5:**
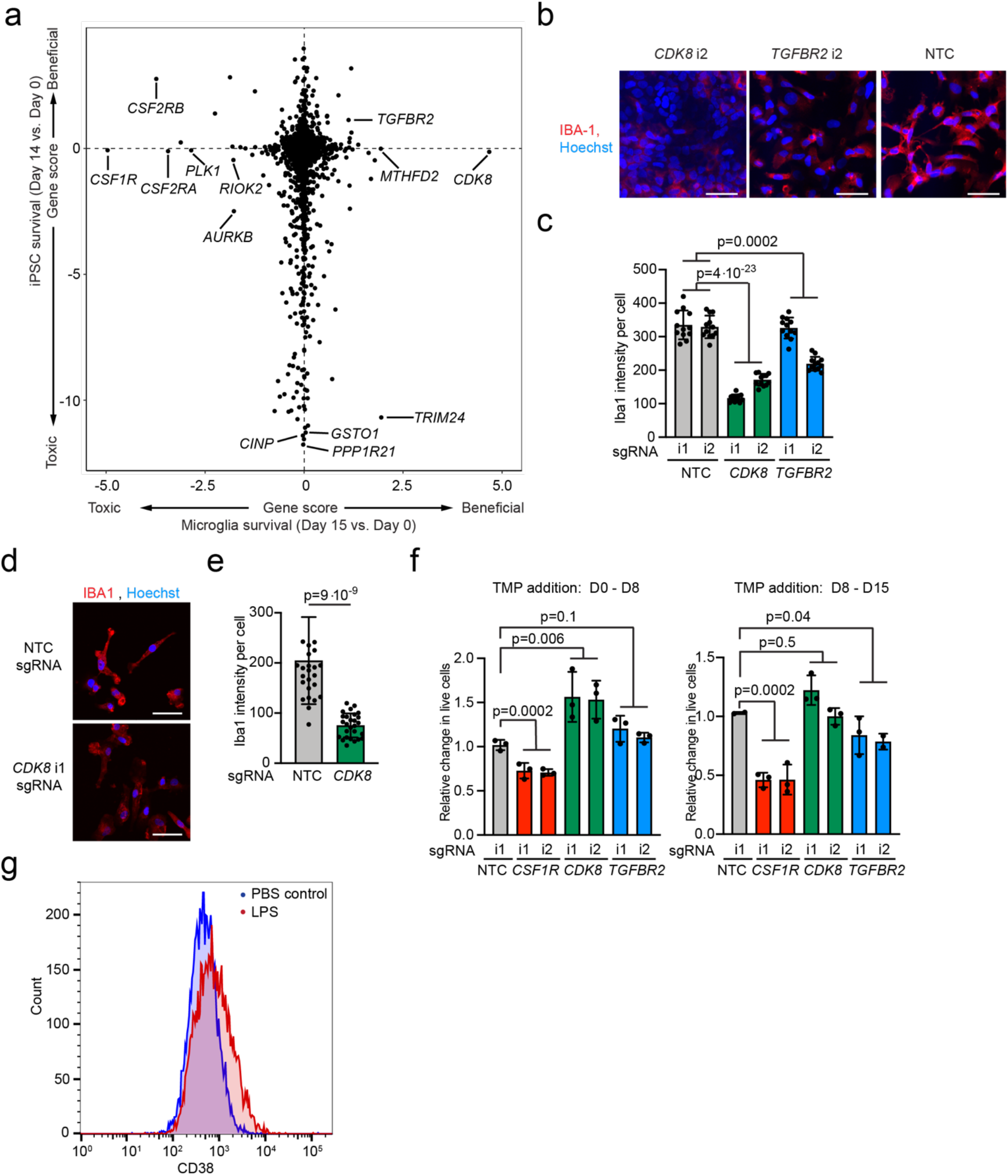
Knockdown of *CDK8* and *TGFBR2* induces proliferation and decreases microglia markers in iPSC-derived microglia generated with different protocols. **a,** Comparison of Gene Scores from CRISPRi survival/proliferation screens in iTF-Microglia (this study) vs. iPSCs^15^. Each dot represents a gene. **b-c**, IBA1 staining in Day 8 CRISPRi iTF-Microglia containing sgRNAs targeting *CDK8* or *TGFBR2* compared to non-targeting control (NTC) sgRNAs. **b**, Representative images. Scale bar = 50 μm. **c**, Quantification. Mean +/-sd, n = 6 fields of view from 2 different wells per sgRNA; p values from two-tailed Student’s t-test. **d-e**, IBA1 staining in Day 8 iMGs generated by the protocol from Brownjohn *et al*., 2018 expressing sgRNAs targeting *CDK8* compared to non-targeting control (NTC) sgRNAs. **d**, Representative images. Scale bar = 50 μm. **e**, Quantification. Mean +/-sd, n =9 fields of view from 3 different wells per sgRNA; p values from two-tailed Student’s t-test. **f**, Relative change in live cells of iTF-Microglia at Day 8 (*left*) and Day 15 (*right*) containing sgRNAs targeting *CDK8*, *CSF1R* or *TGFBR2* compared to non-targeting control sgRNAs. The inducible CRISPRi system was stabilized with TMP from Day 0 – Day 8 (*left*) or Day 8 – Day 15 (*right*). (mean +/-sd, n = 3 biological replicates; p values from two-tailed Student’s t-test. **g,** CD38 cell surface levels measured by flow cytometry in iTF-Microglia 24 h treatement with 100 ng/mL LPS or PBS control.

**Extended Data Figure 6:**
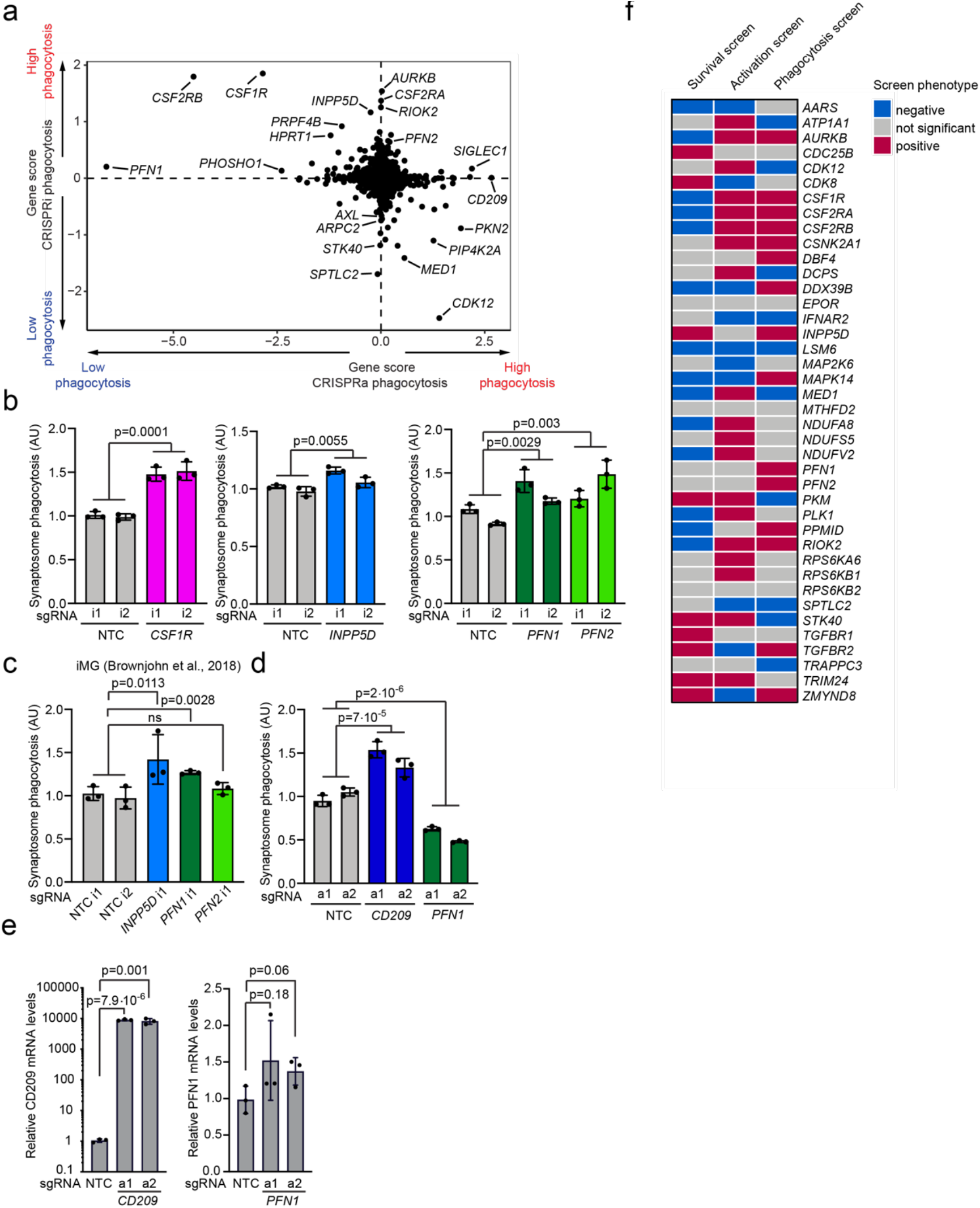
Validation of phagocytosis hits and overview of genes selected for the CROP-seq screen based on primary screens. **a**, Comparing Gene Scores for hits from phagocytosis CRISPRi and CRISPRa screens. Each dot represents a gene. **b-d**, Validation of (b,c) CRISPRi hits and (d) CRISPRa hits in (b,d) iTF-Microglia or (c) iPSC-derived microglia differentiated using an alternative protocol by Brownjohn *et al*.^22^ Phagocytosis of pHrodo-labelled synaptosomes by cells expressing either non-targeting control (NTC) sgRNAs or sgRNAs targeting *CSF1R*, *INPP5D*, *PFN1* and *PFN2* was quantified by flow cytometry. Values represent mean +/- sd of n = 3 biological replicates; p values from two-tailed Student’s t-test. **e**, Overexpression of CD209 (*left*) and PFN1 (*right*) with the inducible CRISPRa system in iTF-Microglia. QPCR quantification of the relative fold change of CD209 and PFN1 mRNA levels in iTF-Microglia expressing CD209 and PFN1 sgRNA as compared to a non-targeting control sgRNA in the presence of TMP (mean +/- sd, n = 3 biological replicates; p values from two-tailed Student’s t-test). CD209 and PFN1 levels were normalized to the housekeeping gene GAPDH. **f**, Binary heatmap of genes selected for the CROP-seq screen and their knockdown phenotype in the CRISPRi survival, phagocytosis and inflammation screens. Red: KD increases phenotype (positive hit). Blue: KD decreases phenotype (negative hit). Grey: not a significant hit, p > 0.1.

**Extended Data Figure 7:**
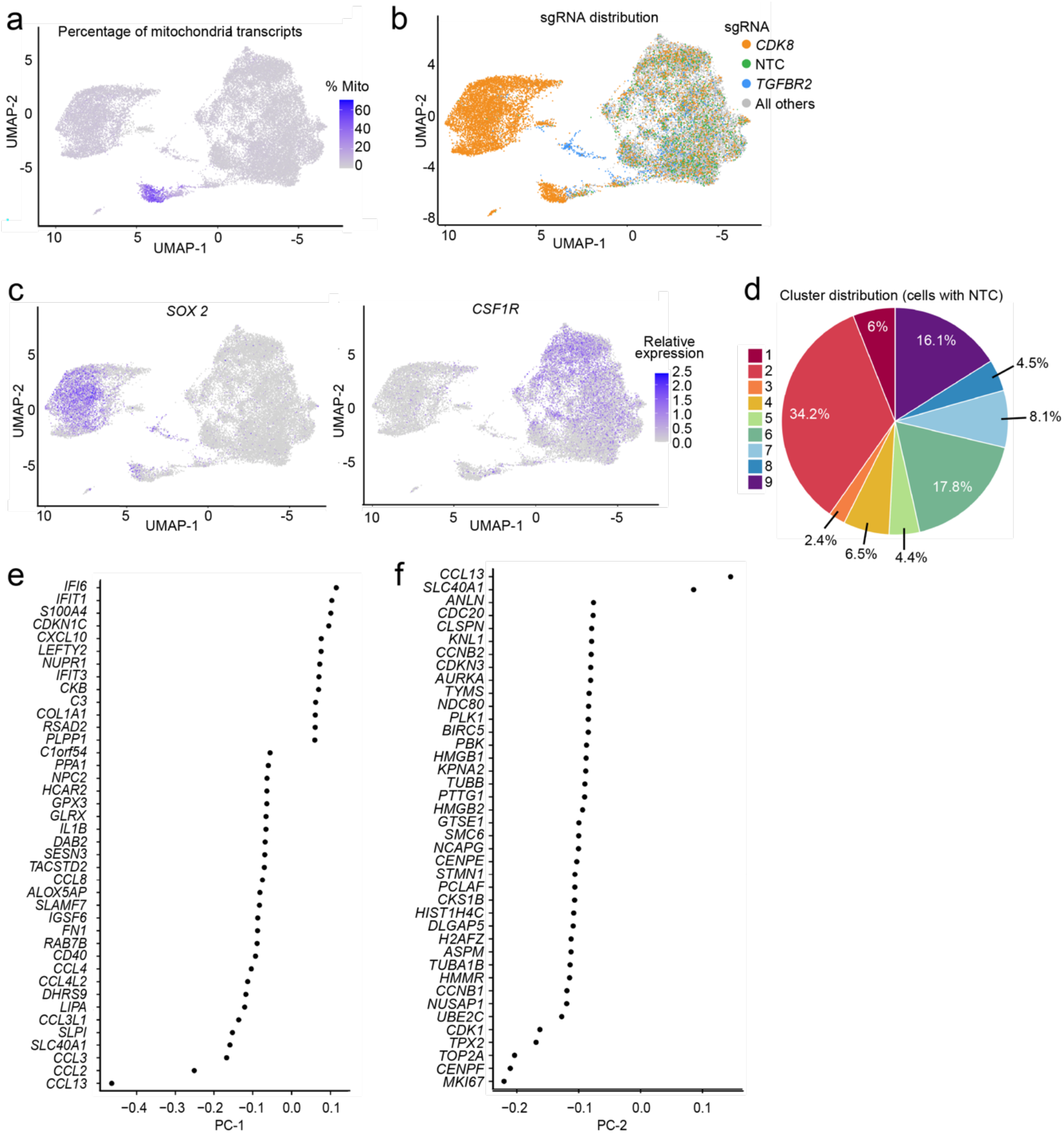
Characterization of microglia cluster signatures. **a-c**, UMAP projection representing single-cell transcriptomes, with cells colored based on (a) the percentage of mitochondrial transcripts, (b) the expressed sgRNAs, with sgRNAs targeting *CDK8* in orange, sgRNAs targeting *TGFBR2* in blue, non-targeting control sgRNAs (NTC) in green, and all other sgRNAs in grey, or (c) expression levels of *SOX2* (*Left*) or *CSF1R* (*Right*). **d**, Distribution of iTF-Microglia expressing non-targeting (NTC) sgRNAs across the 9 clusters described in Figure 6. **e-f**,the top 40 genes with the highest embedding values for (e) the first principal component (PC-1) and (f) the second principal component (PC-2), displayed in ranking order.

**Extended Data Figure 8:**
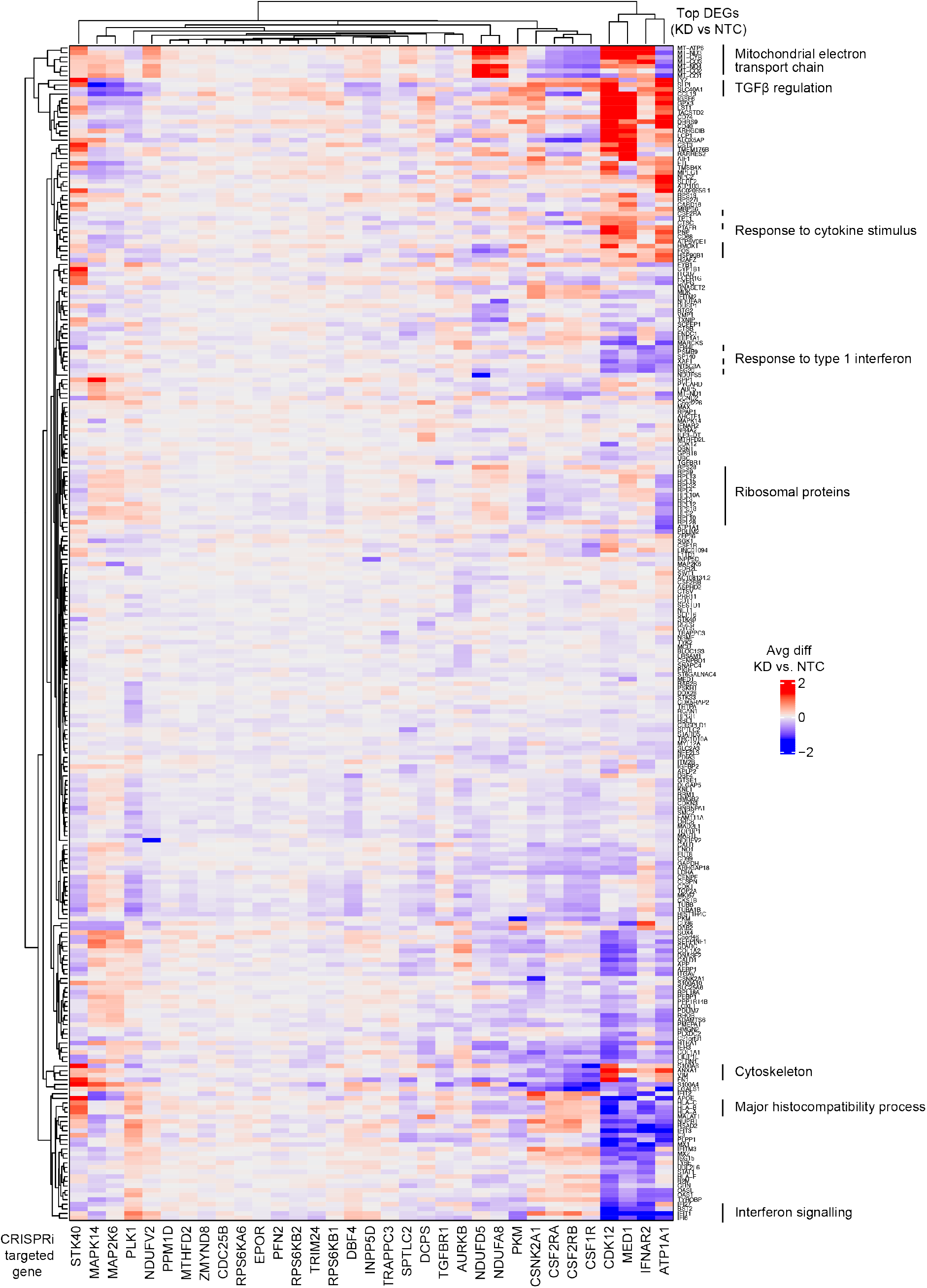
CROP-seq reveals transcriptomic changes in iTF-Microglia induced by gene knockdown. Changes in gene expression in response to CRISPRi knockdown of genes of interest in iTF-Microglia. Each column represents one CRISPRi-targeted gene. For each CRISPRi-targeted gene, cells with the strongest knockdown were selected and the top 20 differentially expressed genes in comparison to non-targeting control (NTC) sgRNA containing cells were selected. The merged set of these genes is represented by the rows. Rows and columns were clustered hierarchically based on Pearson correlation. Functionally related clusters of differentially expressed genes are labeled.

**Extended Data Figure 9:**
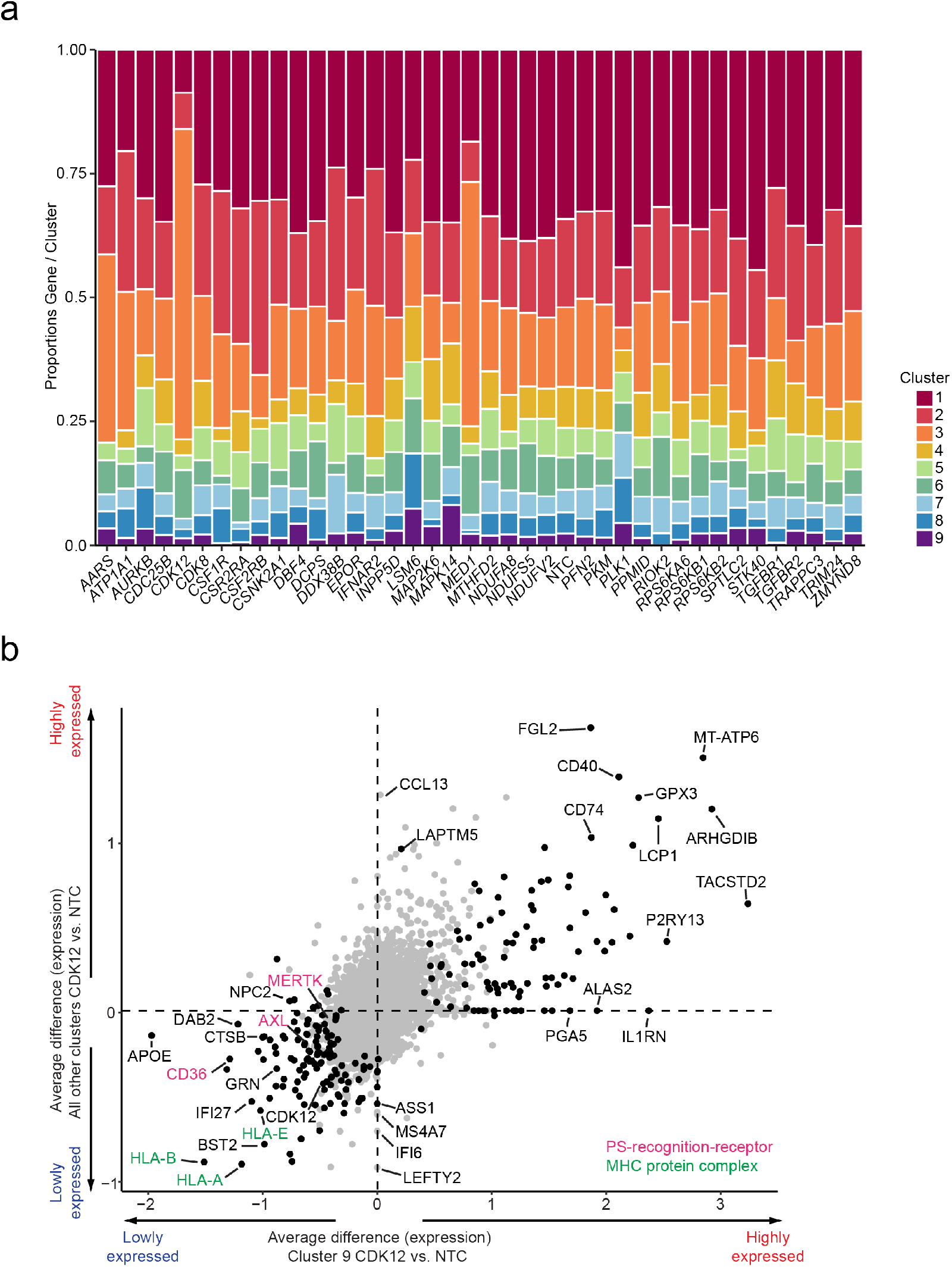
Transcriptomic changes in iTF-Microglia induced by *CDK12* knockdown in cluster 9 and in all other clusters. **a**, Changes in cluster distribution after CRISPRi knockdown of targeted genes in iTF-Microglia. Distribution of cells according to the 37 targeted genes and non-targeting control (NTC) in clusters 1-9. **b**, Average differences of gene expression induced by *CDK12* knockdown in cluster 9 compared to those in all other clusters. Genes encoding phosphatidylserine (PS) recognition receptors are labeled in magenta and Genes encoding MHC complex components are labeled in green.

## SUPPLEMENTARY TABLE LEGENDS

**Supplementary Table 1. RNA-Seq Normalized Counts, Related to Figure 1.** Gene-level counts per sample normalized to library size (transcript per million). Samples, in triplicate, include Day 0 iTF-iPSCs, Day 9 iTF-Microglia (PBS-treated and LPS-treated), Day 15 iTF-Microglia, and Day 9 Brownjohn-iMG (PBS-treated and LPS-treated). Columns are: Ensemble gene ID (ensembl_id), gene, all samples.

**Supplementary Table 2. RNA-Seq LPS Differentially Expressed Genes, Related to Figure 2.** Differentially expressed genes from comparing expression levels of LPS-treated cells to PBS-treated cells in Day 15 iTF-Microglia (first tab) and Brownjohn-iMG (second tab). Columns are: Ensembl gene ID (ensembl id), differentially expressed gene (gene), average expression over all samples (base mean), effect size estimate PBS vs. LPS (log2 fold change), log2 fold change standard error, p value, and adjusted p value. Tab 1 is iTF-Microglia, tab 2 is Brownjohn-iMG.

**Supplementary Table 3. Primary Screen Phenotypes, Related to Figures 4, 5, and Extended Data Figure 6.** Phenotypes from survival and FACS-based screens (survival, activation, and phagocytosis) are listed for all genes targeted in the H1 library. Columns are: targeted transcription start site (index), targeted gene (gene), knockdown phenotype, p value, and the gene score (product of phenotype –log10(p value)).

**Supplementary Table 4. RNA-Seq, Differentially Expressed Genes as a result of *PFN1* overexpression in iTF-Microglia, Related to Figure 5.** Differentially expressed genes of Day 8 iTF-Microglia overexpressing two different *PFN1* sgRNAs compared to non-targeting control sgRNA (NTC).

**Supplementary Table 5. CROP-seq Pooled sgRNA Library, Related to Figures 6 and 7.** Sequences for sgRNAs in CROP-seq pooled sgRNA library. Columns are: gene targeted for CRISPRi knockdown (target.gene), sgRNA short name as used in the paper (sgRNA.name), and sgRNA protospacer sequence (sgRNA.sequence).

**Supplementary Table 6. Overview of CROP-seq results, Related to Figures 6 and 7.**

**Supplementary Table 7. CROP-seq Cluster Differentially Expressed Genes, Related to Figure 6.** Differentially expressed genes (DEGs) for each UMAP cluster (1-9) compared to all other cluster, only positive values included. Columns are: gene, cluster, average log2 fold change, p value, adjusted p value

**Supplementary Table 8. CROP-seq Target Gene Cluster Proportions, Related to Figure 7.** Relative proportions of cells in clusters (rows) with a sgRNA targeting a given gene (columns) normalized to non-targeting control proportions (NTC).

**Supplementary Table 9. CROP-seq Knockdown versus Control Differentially Expressed Genes, Related to Extended Data Figure 9.** Differentially expressed genes, calculated using Student’s T-test between cell with CRISPRi knockdown and non-targeting control (NTC) sgRNAs. Columns are: gene targeted for CRISPRi knockdown (TargetGene), differentially expressed gene (Gene), log2 counts per million, log2-fold change, p value, false discovery rate (FDR).

**Supplementary Table 10. Individual sgRNA sequences and primers.** qPCR and CROP-seq sgRNA enrichment primers, as well as individually cloned sgRNAs are listed. Columns are: name of sgRNA or primer (Name), section of sequence (sequence), sequence 5’ to 3’, use in this study (use).

