## Supplementary Figure 1 for "A CRISPRi/a platform in iPSC-derived microglia uncovers regulators of disease states"

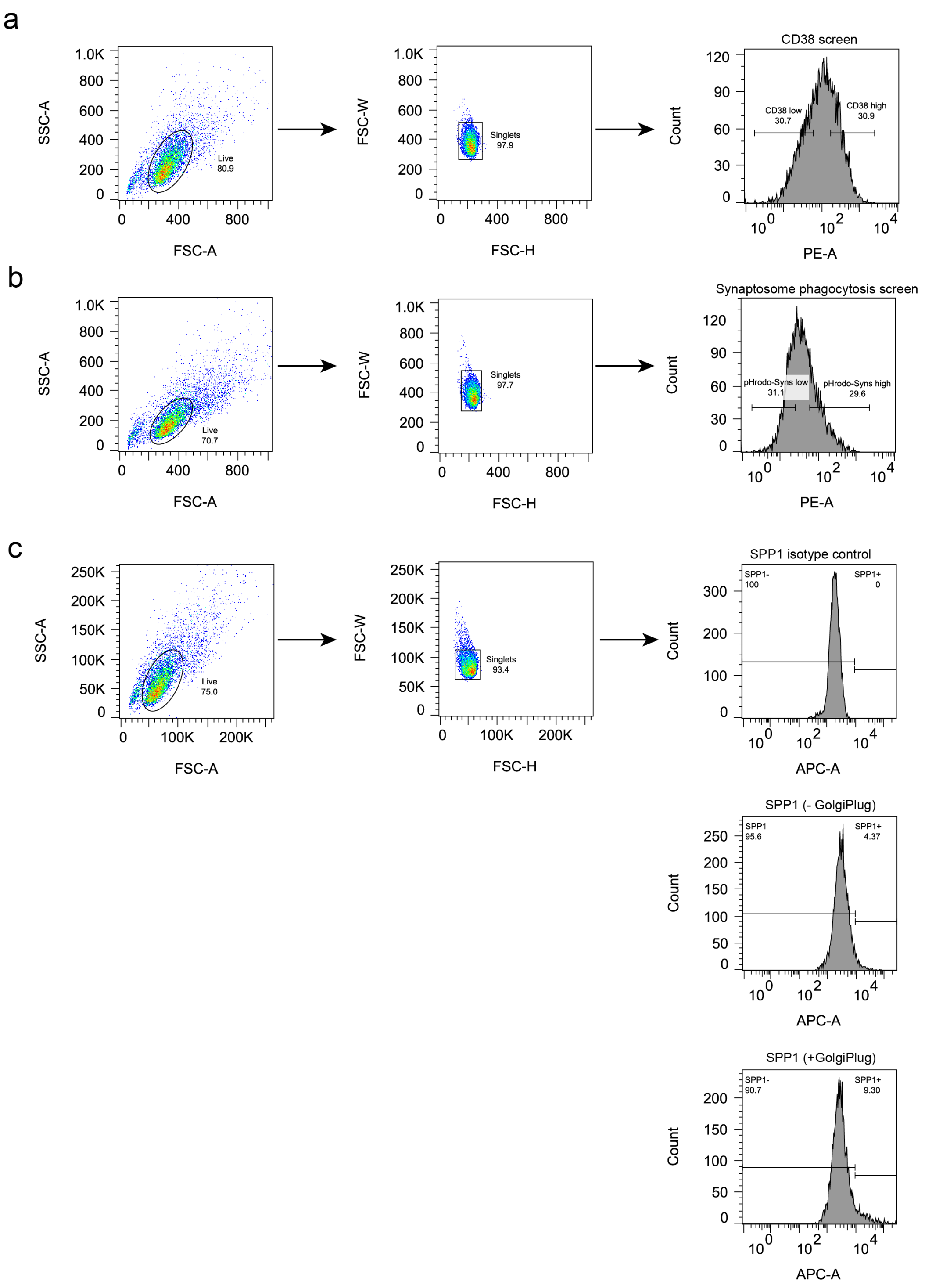


**Supplementary Figure 1: Gating strategies for FACS-based screens and SPP1 positive cells. a,b** Gating strategies for (a) CD38 screens and (b) synaptosome phagocytosis screens. Intact iTF-Microglia were identified from FSC-SSC plot and then gated for singlets. These cells were sorted into high and low signal populations corresponding to the top 30% and the bottom 30% of the signal distribution. **c,** To determine the fraction of SPP1+ cells, cells were treated with GolgiPlug and singlets were classified using the SPP1 isotype control to determine the threshold.
